# Cancer-associated fibroblasts produce matrix-bound vesicles that influence endothelial cell function

**DOI:** 10.1101/2023.01.13.523951

**Authors:** Alice Santi, Emily J Kay, Lisa J Neilson, Lynn McGarry, Sergio Lilla, Margaret Mullin, Nikki R Paul, Frédéric Fercoq, Grigorios Koulouras, Giovanny Rodriguez Blanco, Dimitris Athineos, Susan Mason, Mark Hughes, Yann Kieffer, Colin Nixon, Karen Blyth, Fatima Mechta-Grigoriou, Leo M Carlin, Sara Zanivan

## Abstract

Intercellular communication between different cell types in solid tumors contributes to tumor growth and metastatic dissemination. The secretome of cancer-associated fibroblasts (CAFs) plays major roles in these processes. Using human mammary CAFs, we unveil a mechanism of cell-cell communication between CAFs with myofibroblast phenotype and endothelial cells (ECs) based on intercellular protein transfer through extracellular vesicles (EVs). CAFs transfer proteins to ECs, including plasma membrane receptors, which we have identified by using mass spectrometry- based proteomics. Using THY1 as an example of transferred plasma membrane-bound protein, we show that CAF-derived proteins can influence how ECs interact with other cell types. Here, we show that CAFs produce high amounts of matrix-bound EVs that have a key role in protein transfer. Hence, our work paves the way for further studies to understand how CAF-derived matrix-bound EVs influence tumor pathology by regulating functions of neighboring cancer, stromal and immune cells.

**One sentence summary:** CAFs with a myofibroblastic-like phenotype transfer proteins to ECs, including plasma membrane receptors, through matrix-bound EVs

## Introduction

The communication between cells is fundamental for the physiological function of tissues (1, 2) and alterations can cause diseases and determine their severity (3–5). In solid tumors, intercellular communication involves cancer cells and neighboring cells of the tumor microenvironment (TME) and modulates tumor growth and metastatic dissemination. The TME is a highly heterogeneous and dynamic compartment that comprises of pathologically and activated immune and stromal cells, which include cancer-associated fibroblasts (CAFs) and endothelial cells (ECs) (6, 7).

CAFs are highly secretory cells and represent the bulk of the stroma of solid tumors with a desmoplastic reaction, such as breast cancer (BC) (8, 9), thus being a considerable source of chemical signals that can affect the behavior of cancer, immune and stromal cells. For these reasons, CAFs have been defined as “architects of cancer pathogenesis” (10) or as “architects of stroma remodeling” (6). The repertoire of chemical signals produced by CAFs includes growth factors, cytokines, non-coding RNAs, components of the extracellular matrix (ECM) and ECM remodeling enzymes, which regulate invasion, proliferation and chemoresistance of cancer cells, blood vessel formation and the recruitment and function of immune cells (6, 10–14). It is now evident that CAFs carry out these different functions by acquiring distinct but interchangeable states (15). Myofibroblast-like CAFs (myCAFs) and inflammatory CAFs (iCAFs) are the two main subtypes that have been described in tumors, including BC (15, 16). myCAFs are responsible for ECM production and remodeling and have immunosuppressive functions, while iCAFs have an immunomodulatory role (15, 17). In addition to these classical mechanisms of paracrine crosstalk, CAFs transfer a wide range of nutrients (18–20), proteins, lipids (21, 22) and even entire mitochondria (23, 24) to cancer cells, which use these CAF-derived resources to support their own growth and motility.

The intercellular transfer of cell surface and intracellular proteins has been extensively documented between immune cells. The physiological role and functional consequences of this phenomenon are still unclear, but it has been suggested that it may help to regulate the immune response (25–29). So far, few works have examined the ability of pathologically activated fibroblasts to transfer their own proteins to cancer cells. These works showed that the transfer of proteins from CAFs to cancer cells occurs through large extracellular vesicles (EVs) that CAFs release in the conditioned medium (CM) and that it supports cancer cell proliferation (21) and migration (22). The fact that only a small number of studies have investigated the protein transfer ability of CAFs leaves still many open questions: do stromal and immune cells also receive CAF-derived proteins? If so, what is the biological relevance of this intercellular protein transfer?

EVs are lipid bilayer-enclosed particles that mediate cell-cell communication by transferring proteins, lipids and nucleic acids between cells. In accordance with the MISEV guidelines, EVs are classified based on their size as small (diameter < 100-200 nm) and medium/large (diameter > 150-220 nm) (30). Medium/large EVs directly bud from the plasma membrane (ectosomes), while small EVs originate from either the endosomal compartment (exosomes) or the plasma membrane (ectosomes) (30, 31). EVs that transfer biological material between cells are typically found in cell-derived CM (CM-EVs) (14); however, few studies have demonstrated the presence of EVs embedded within the ECM of decellularized tissues and of murine NIH-3T3 fibroblast cell cultures (32, 33). These matrix-bound vesicles (MBVs) have a similar shape and morphology to CM-EVs, but differ in lipid and microRNA content (33). MBVs are biologically active (32, 34); however their protein composition and role in intercellular protein transfer have not yet been reported.

Tumor blood vessels are typically embedded within the tumor stroma, therefore, we have investigated whether CAFs employ intercellular protein transfer to influence the function of ECs. Using CAFs isolated from patients with BC as donors and human ECs as recipient cells, we have identified a specific pool of proteins that CAFs transfer to ECs and, using THY1 as example, we provide a proof of principle that they can be functional in the ECs. Moreover, we found that CAFs deliver proteins principally through MBVs and that CAFs expressing myCAF markers are the main donors of proteins to ECs.

## Results

### CAFs transfer proteins to ECs

To study whether mammary CAFs transfer proteins to ECs, we used several CAF lines that we have isolated from patients with BC (pCAFs). These pCAFs express the mesenchymal marker vimentin (Fig. S1A) (35), but are negative for markers of epithelial, endothelial and immune cells (Fig. S1B). Our lab has previously characterized the lines pCAF2 and pCAF3 (35). To monitor the transfer of proteins from pCAFs (donor cells) to human umbilical vein ECs (HUVECs, recipient cells), we fluorescently labeled the pCAF proteome with CFSE, a dye that covalently binds to amino groups. Microscopy analysis showed that HUVECs became fluorescent after being co-cultured for 24h with CFSE-labeled pCAFs, indicating that pCAFs transferred some of their proteins to HUVECs (Fig. 1A and Fig. S1C).

**Fig. 1.**
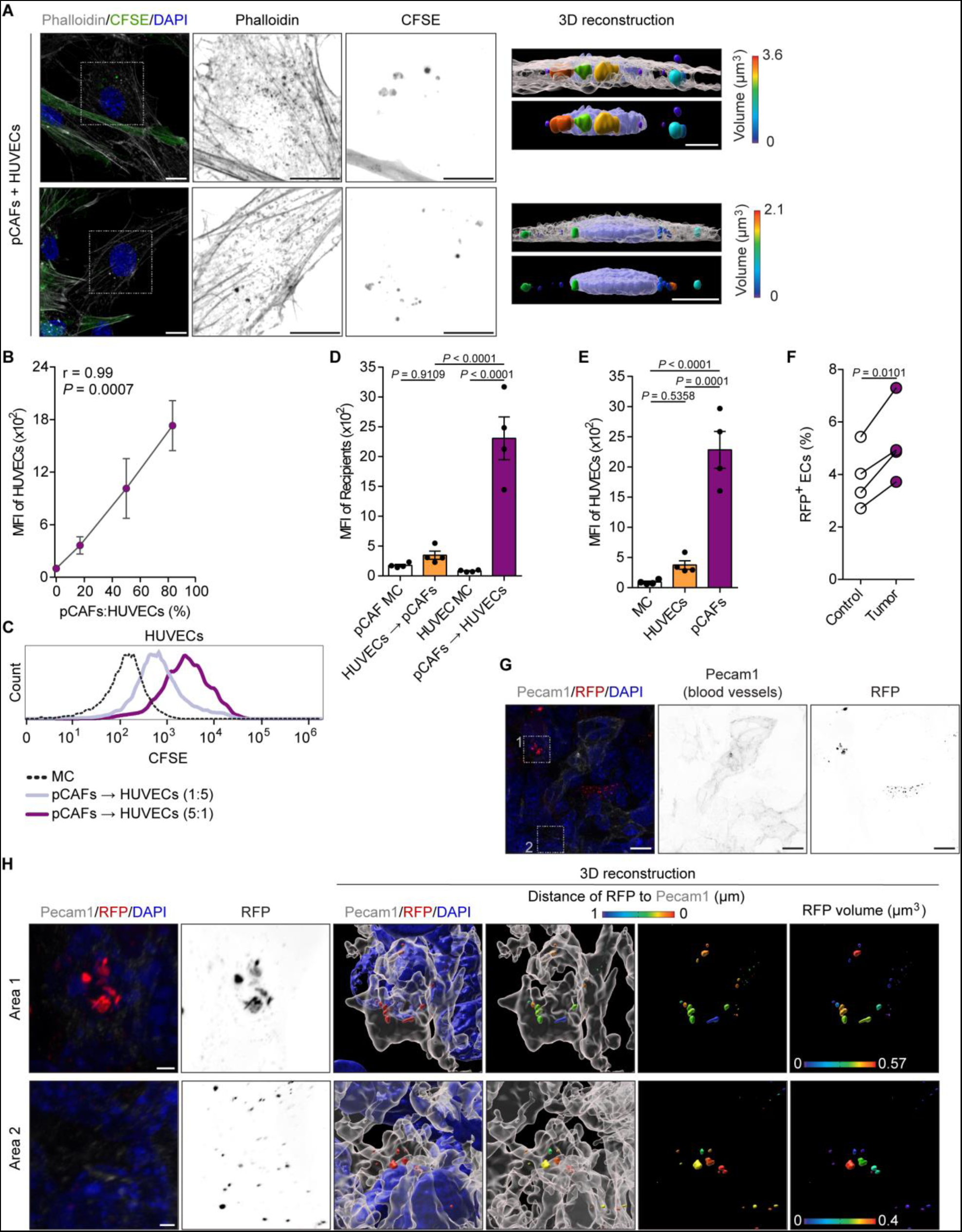
CAFs transfer proteins to ECs in vitro and in vivo. **(A)** Representative images (maximum intensity projection processing from confocal z-stack) and corresponding 3D reconstruction of protein transfer from CFSE-labeled pCAFs (fully green cells) to HUVECs. Actin and nuclei were stained with phalloidin and DAPI, respectively (scale bar = 10 μm). **(B, C)** Quantification **(B)** of the protein transfer from pCAFs to HUVECs by altering the ratio between the two cell types, pCAFs:HUVECs ratios are 1:5, 1:1, 5:1, *N =* 3 (MFI, median fluorescence intensity and r, Pearson correlation) and representative histogram **(C)** of the 1:5 and 5:1 ratio (the y-axis is normalized to mode). **(D, E)** Comparison between “HUVECs to pCAFs” **(D)** or “HUVECs to HUVECs” **(E)** and “pCAFs to HUVECs” protein transfer ability, *N =* 4 (MC, monoculture). **(F)** Proportions of RFP^+^ ECs in lungs of α-SMA-RFP tumor-free mice or mice with lung metastases, *N =* 4 mice/condition, paired mice were born on the same day. **(G)** Representative image (maximum intensity projection processing from confocal z-stack) of the tumor area in the lung of α-SMA-RFP mice stained for Pecam1. Nuclei were stained with DAPI, scale bar = 10 μm. **(H)** 3D reconstruction of the tumor vasculature and of the RFP signal, the distance between the RFP signal and the endothelium and the volume of the RFP signal are shown, scale bar = 2 μm. **(D, E)** Data are represented as mean ± SEM, one-way ANOVA with Tukey’s multiple comparison test. **(F)** Two-tailed paired t test.

Using the same CFSE-based labeling method, we quantified the intercellular transfer of proteins by flow cytometry, which confirmed that HUVECs acquire fluorescent signals upon co-culture with CAFs (Fig. 1B-E). Notably, the quantity of transferred proteins was dependent on the number of donor cells and it increased in accordance with the ratio between pCAFs and ECs (Fig. 1B-C). The shift of the CFSE peak of co-cultured HUVECs compared with monoculture showed that the vast majority of the HUVECs received pCAF proteins, indicating that this is a commonly occurring event (Fig. 1C). Conversely, HUVECs transferred very low amounts of proteins to pCAFs (Fig. 1D) or to HUVECs themselves (Fig. 1E). These results indicate that pCAFs and HUVECs do not mutually exchange proteins and that CAFs are the major protein donors.

Once we established that pCAFs transfer proteins to HUVECs in vitro, we sought to assess whether this mechanism also occurred in vivo. To this purpose, we used the C.FVB-tg(Acta2-DsRed)1RK1/J mouse model (36), also known as α-SMA-RFP. This model expressed the red fluorescent protein(RFP) in cells expressing the alpha-smooth muscle actin gene (*Acta2,* whose product is α-SMA protein). Because α-SMA is a widely used CAF marker (7, 15, 37), we used the α-SMA-RFP model to monitor the transfer of RFP from *Acta2* expressing cells to ECs in experimental pulmonary metastases, as a mean of protein transfer from CAFs to ECs. 4T1 murine BC cells were injected in the tail vein of α-SMA-RFP mice and, after three weeks, we dissected the lungs and analyzed single cell suspension by flow cytometry. We used α-SMA-RFP mice that had not been injected with 4T1 cells as control to measure whether any RFP protein could be transferred to the endothelium in the absence of *Acta2* expressing CAFs (e.g. by perivascular cells, such as pericytes, which also express *Acta2*). Flow cytometry analysis measured a significant increase of RFP^+^ ECs in mice with lung metastases compared with the control (Fig. 1F). To confirm these results, we imaged fixed precision cut lung slices with 4T1 metastases from α-SMA-RFP mice (Fig. 1G). The 3D reconstruction of tumor sections, which were stained for platelet endothelial cell adhesion molecule (Pecam1) to visualize ECs, showed RFP^+^ endothelium in the lung metastases of these mice (Fig. 1G-H), but not in non-RFP expressing control mice (Fig. S1D).

Overall, our data provide evidences that CAFs communicate with ECs through the transfer of proteins in vitro and possibly also in vivo.

### CAFs transfer plasma membrane receptors to ECs

To identify proteins that pCAFs transfer to HUVECs, we used a mass spectrometry (MS)-based trans-stable-isotope labeling of amino acids in cell culture (trans-SILAC) proteomic approach (28). First, we labeled the proteome of pCAFs with the heavy isotopologue of arginine and lysine, and co-cultured them with unlabeled HUVECs for 4h or 24h. Then, we sorted the HUVECs and analyzed their proteome by MS (Fig. 2A). We quantified 808 and 1062 heavy-labeled proteins in at least three out of five biological replicates at 4h and 24h time point, respectively (Fig. 2B and Data S1). Of these, 698 proteins were common to both time points (Fig. 2B). Gene Ontology Cellular Component (GOCC) term analysis of the proteins transferred from CAFs to the HUVECs revealed enrichment in lipid bilayer-enclosed vesicles, endoplasmic reticulum (ER), ER-Golgi intermediate compartment, and macromolecular complexes, including focal adhesions, cell junctions, ribonucleoprotein particles and proteasome (Fig. 2C). The high number of common proteins and the consistency of the top ten enriched GO terms between the two time points indicates that, in culture, there is a continuous transfer of proteins over time from CAFs to ECs. Moreover, the association of these proteins to particular subcellular compartments suggests that mammary CAFs transfer selected protein subsets.

**Fig. 2.**
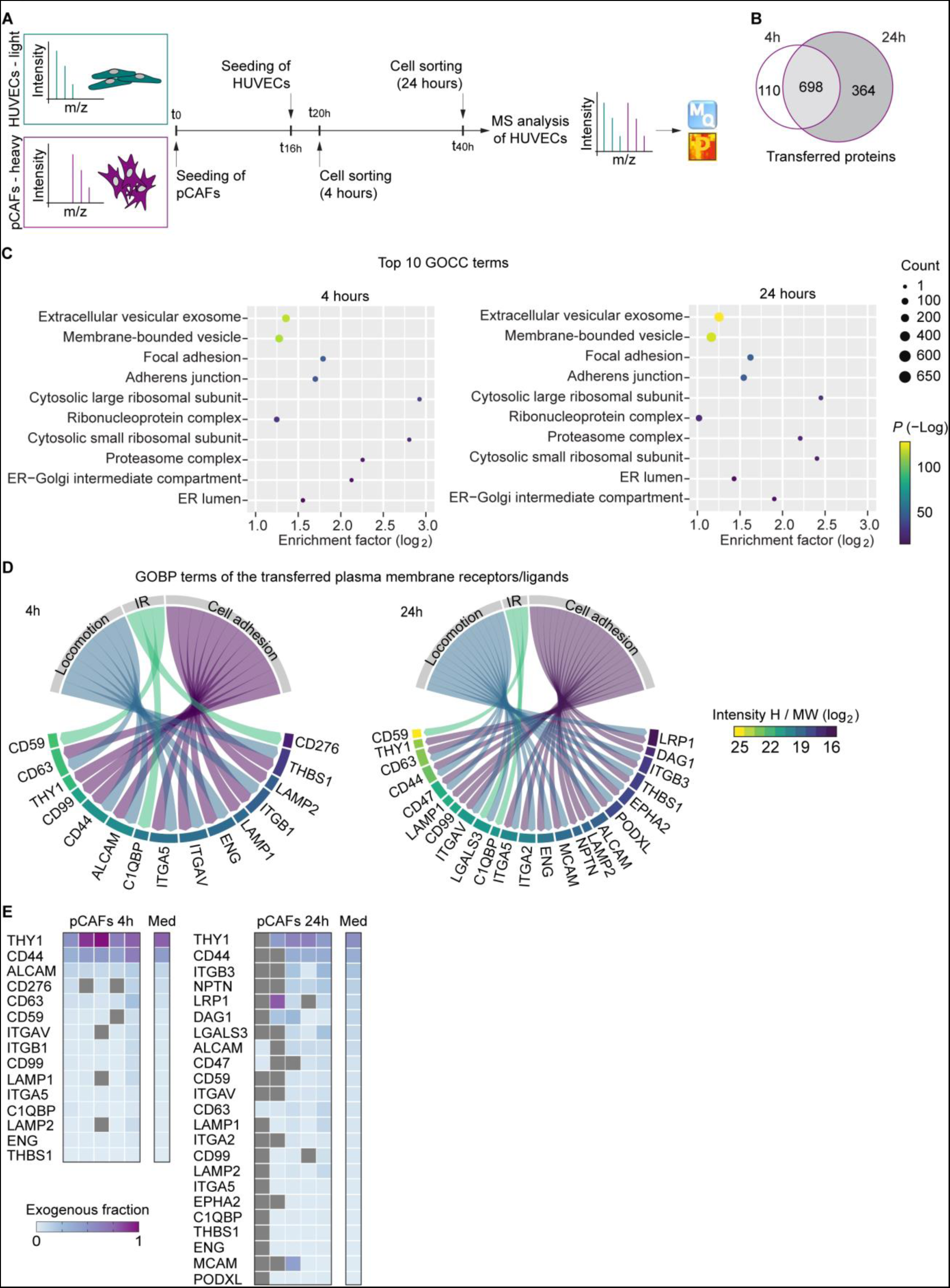
Identification of the proteins transferred from CAFs to ECs. **(A)** Workflow diagram of the trans-SILAC experiment for the identification of the proteins transferred from pCAFs to HUVECs. **(B)** Venn diagram showing the number of transferred proteins at each time point. **(C)** Top 10 GOCC terms based on descending *P* (–Log) and with at least a two-fold enrichment. The enrichment analysis of the transferred proteins was performed using pCAF proteome (Data S6) as reference (Fisher exact test). **(D)** Classification of the plasma membrane receptors/ligands based on the GO biological process (GOBP) terms. Proteins are sorted by decreasing values of the median of the intensity value in the “heavy” channel of the trans-SILAC experiment (Intensity H) divided by the MW in the log_2_ scale, *N =* 5 (IR, immune response). **(E)** Heatmap showing the exogenous fraction of the transferred receptors/ligands for each independent experiment and their median. Proteins are sorted by decreasing values of the median of the exogenous fraction.

Previous studies have shown that cancer and immune cells use EVs to transfer functional plasma membrane proteins to ECs (38, 39). These types of proteins are highly relevant because they may alter the function of the endothelium, including its interactions with surrounding cells. Therefore, we focused our analysis on plasma membrane receptors and membrane-bound ligands. Interestingly, the majority of the transferred membrane proteins were involved in immune response, cell locomotion, cell-cell and cell-matrix adhesion (Fig. 2D) (40–44), corroborating the idea that CAF-derived proteins may have important implications on the functions of the tumor vasculature. To select for proteins that provided the biggest changes in the HUVEC proteome, we determined the contribution of each transferred protein to the corresponding endogenous protein in the HUVECs and referred to this value as “exogenous fraction”. The exogenous fraction ranges between 0 and 1, and the closer the value is to 1, the more the pCAF protein contributes to the endothelial counterpart (Fig. 2E and Data S1). CAF-derived Thy-1 membrane glycoprotein (THY1) was the protein with the highest contribution to the HUVEC proteome, with an exogenous fraction of 0.78 and 0.54 after 4h and 24h of co-culture, respectively (Fig. 2E and Data S1). CD44 antigen (CD44) also contributed highly with an exogenous fraction of 0.46 and 0.35 at 4h and 24h, respectively, and then integrin beta-3 (ITGB3), with an exogenous fraction of 0.21 at 24h of co- culture. The exogenous fraction for all the other receptors and ligands was lower than 0.15 (Fig. 2E and Data S1). Overall, these results indicate that mammary CAF-derived receptors/ligands can quantitatively modify the proteome of the HUVECs. Next, we sought to determine whether transferred proteins could functionally change the HUVECs and focused on THY1, since it contributed to a major change of the HUVEC proteome.

### CAF-derived THY1 induces functional changes in ECs

To confirm that the THY1 detected in HUVECs was derived from pCAFs, rather than being expressed by HUVECs when co-cultured with them, we measured *THY1* transcript in HUVECs in monoculture and after 24h of co-culture with pCAFs. Using pCAFs as control of *THY1* expressing cells, we found that *THY1* mRNA levels did not increase significantly in co-cultured HUVECs compared to the monoculture (Fig. 3A). In addition, flow cytometry analysis confirmed the transfer of THY1 from pCAFs to HUVECs (Fig. 3B-D). While THY1 was not present at the surface of HUVECs in monoculture, after 24h of co-culture with pCAFs, the majority of HUVECs positively stained for THY1 (Fig. 3C-D). Moreover, pCAFs silenced for *THY1* (Fig. 3B) transferred significantly less receptor to the HUVECs (Fig. 3C-D), while the total amount of transferred proteins was not affected (Fig. 3E).

**Fig. 3.**
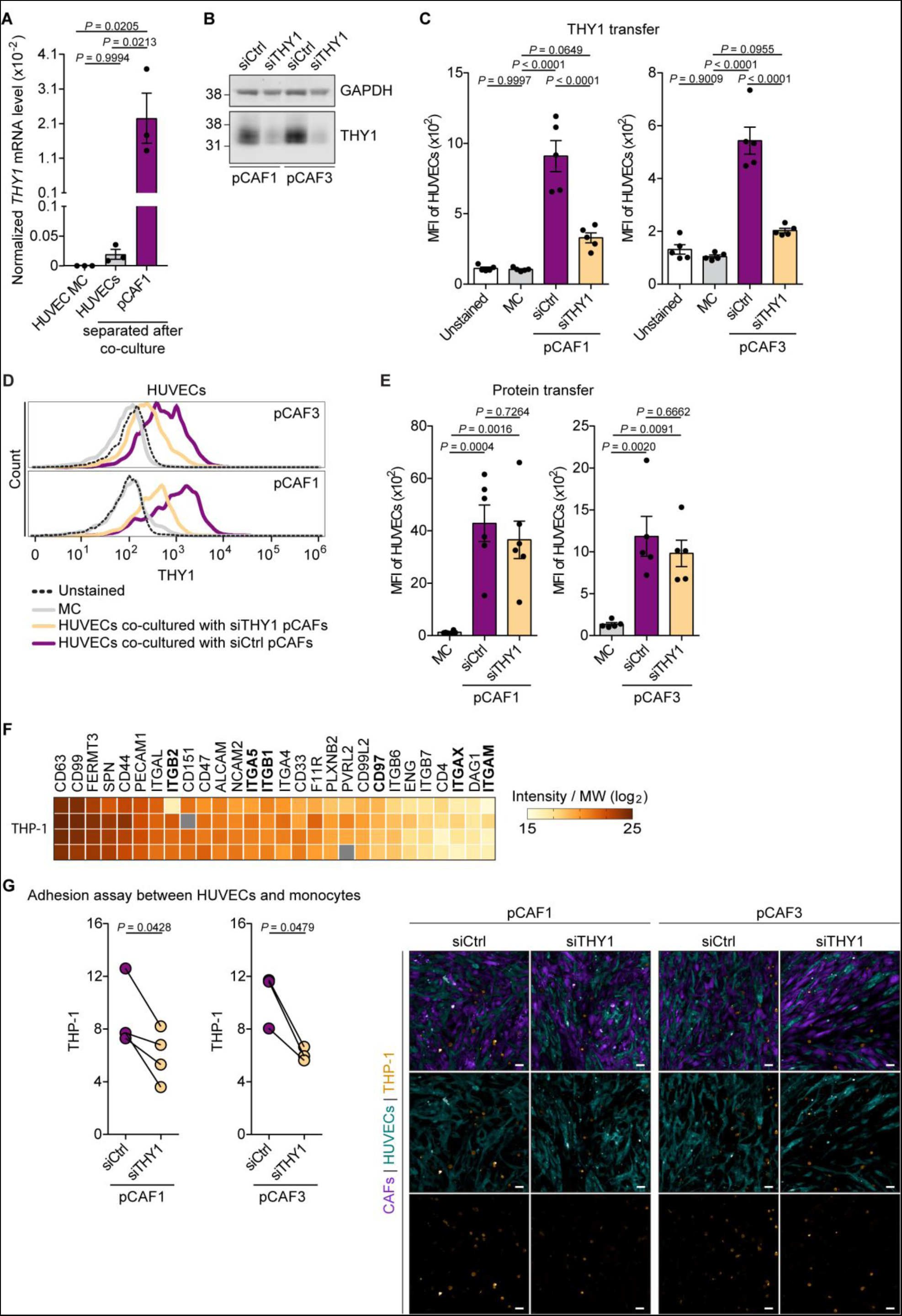
CAF-derived THY1 supports the physical interaction between ECs and monocytes. **(A)** mRNA expression of *THY1* in HUVECs in monoculture and after 24h of co-culture with pCAFs, and in pCAFs. *THY1* mRNA levels were normalized to *18S* expression, *N =* 3. **(B)** Representative western blot showing THY1 protein levels in pCAFs transfected with siCtrl or siTHY1, GAPDH is used as loading control. **(C, D)** Quantification **(C)** of THY1 protein levels in monoculture of HUVECs and HUVECs that were co-cultured with pCAFs transfected with siCtrl or siTHY1 (*N =* 5) and representative histogram **(D)** of THY1 protein levels in HUVECs (the y-axis is normalized to mode). **(E)** Quantification of the protein transfer from pCAFs transfected with siCtrl or siTHY1 to HUVECs, *N =* 6 (pCAF1) and 5 (pCAF3). **(F)** Heatmap based on intensity value divided by the MW in the log_2_ scale of the cell adhesion molecules that were identified in THP-1 monocytes in each independent experiment. THY1 binding partners are in bold. **(G)** Number of THP-1 monocytes per field that bound HUVECs that were co-cultured with pCAFs silenced or not for THY1, *N =* 4 (pCAF1) and 3 (pCAF3), and representative images, scale bar = 50 μm. **(A, C, E)** Dara are represented as mean ± SEM, one-way ANOVA with Tukey’s multiple comparison test, **(G)** Two-tailed paired t test.

THY1 (also known as CD90) is a glycophosphatidylinositol-anchored protein that localizes on the extracellular side of the plasma membrane of cells and that binds to cancer cells and leukocytes through plasma membrane receptors, such as integrin αvβ3, αvβ5, α5β1, αxβ2, αMβ2, syndecan-4, and adhesion G protein-coupled receptor (ADGRE5, also known as CD97) (45, 46). To assess the function of pCAF-derived THY1, we measured leukocyte adhesion to HUVECs when co-cultured with pCAFs silenced or not for THY1. Specifically, we used the human monocyte cell line THP-1 that expresses several THY1 binding partners (Fig. 3F and Data S2). Microscopy analysis of the co-cultures showed that significantly fewer monocytes adhered to the HUVECs when co-cultured with THY1-silenced pCAFs compared with control co-culture (siCtlr), supporting the functionality of THY1 on the HUVEC surface (Fig. 3G). Hence, CAF-derived THY1 endows HUVECs with additional cell-cell adhesion properties.

### MBVs have a major role in protein transfer to ECs

Next, we investigated how pCAFs transfer their proteins to HUVECs. Our data showed that a high number of transferred proteins belonged to lipid bilayer-enclosed vesicles (Fig. 2C), supporting that EVs can be a major route of intercellular protein transfer. It is known that CM-EVs are involved in protein transfer (21, 22, 47–49), but the role of MBVs has not been investigated.

We isolated EVs from both the CM and extracellular matrix of pCAFs (Fig. 4A). Electron microscopy analysis showed that the two EV types had a similar morphology (Fig. 4B). Nanoparticles tracking analysis showed that the diameter of both types of EVs ranged between 50 and 350 nm (Fig. 4C). However, the amount and size distribution differed between the two EV types. Those in the CM mainly included small particles with a diameter between 50 and 150 nm, while MBVs mostly consisted of large EVs, with major peaks at 150 nm and 200 nm (Fig. 4C-D).

**Fig. 4.**
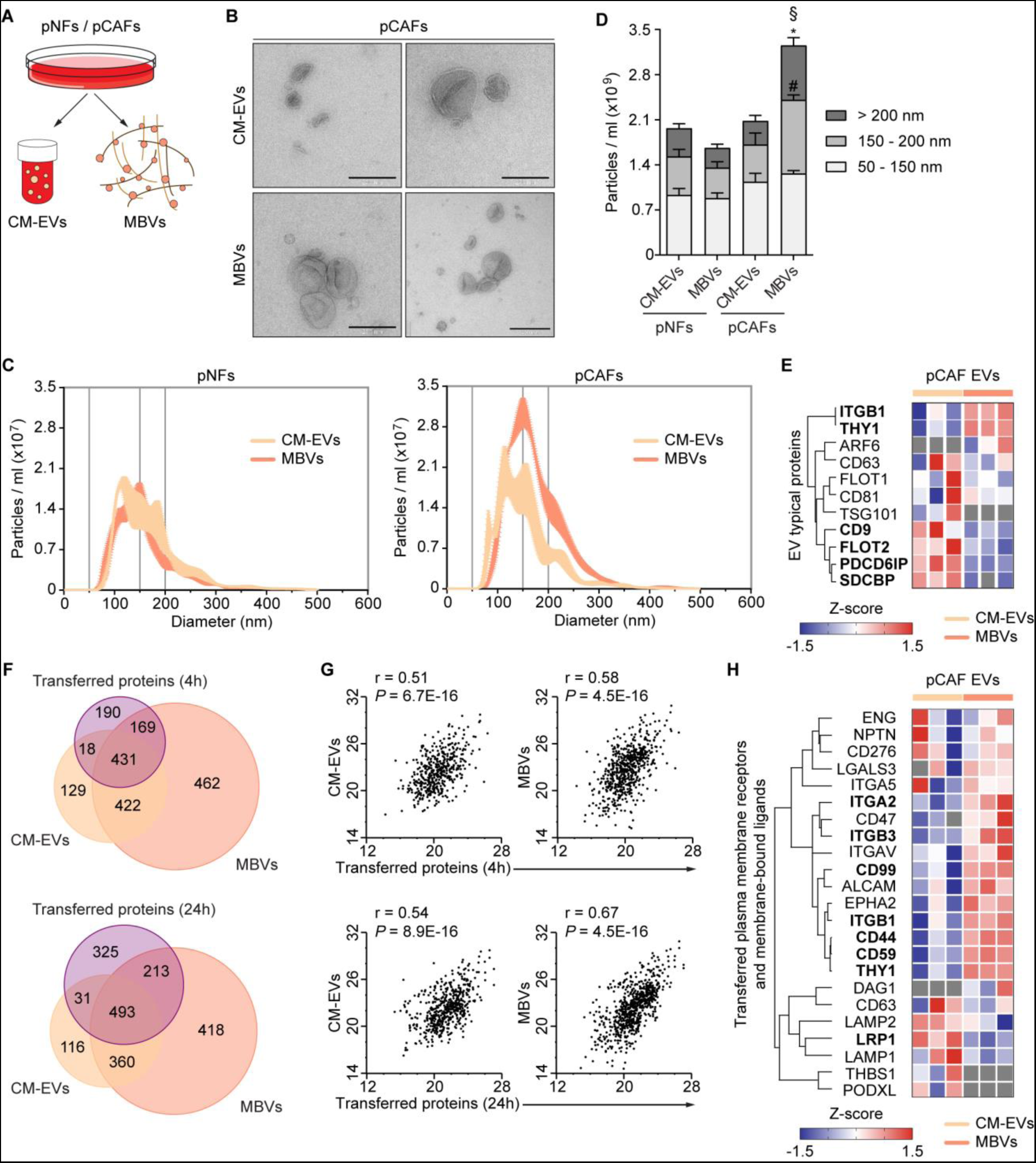
CAF-derived EVs are carriers of the transferred proteins. **(A)** Schematic representation of the purified EV types. **(B)** Representative electron microscopy images of pCAF-derived CM-EVs and MBVs. **(C, D)** Frequency plot **(C)** and histogram **(D)** showing the total amount and size distribution profile of the CM-EVs and MBVs secreted by pNFs and pCAFs using Nanoparticles Tracking Analysis. **(E)** Hierarchical clustering based on average Euclidean distance and heatmap based on the Z-score of the LFQ intensity (log_2_) calculated for the EV typical proteins in the EV proteome, in bold are the proteins significantly different in abundance (two-sided Student’s T-test, *P* < 0.05), *N =* 3/EV type. **(F)** Venn diagram (based on the protein gene names) of the transferred proteins and of the proteins identified by MS proteomics in CM-EVs and MBVs. **(G)** Scatter plot showing the correlation between the amount of the transferred proteins and their relative content in CM-EVs and MBVs. The y- and x-axis report the median of the intensity divided by MW in the log_2_ scale (r, Pearson correlation), *N =* 5 (trans-SILAC experiment) and 3 (for each EV type). **(H)** Hierarchical clustering based on average Euclidean distance and heatmap based on the Z-score of the LFQ intensity (log_2_) calculated for the transferred plasma membrane receptors and membrane-bound ligands in the EV proteome, in bold are the proteins significantly different in abundance (two-sided Student’s T-test, *P* < 0.05), *N =* 3/EV type. **(D)** Data are represented as mean ± SEM, one-way ANOVA with Tukey’s multiple comparison test: * *P* ≤ 0.05, it compares the same EV fraction between pCAF-derived MBVs and pNF-derived CM- EVs or MBVs or pCAF-derived CM-EVs; # *P* ≤ 0.05, it compares the same EV fraction between pCAF-derived MBVs and pNF-derived MBVs or pCAF-derived CM-EVs; § *P* ≤ 0.05, it compares the total amount of pCAF-derived MBVs with respect to pNF-derived CM-EVs or MBVs.

We molecularly characterized pCAF-derived EVs using MS proteomics (Data S3). This analysis confirmed that both types of particles contain common EV markers, such as tetraspanins CD63, CD81, CD9 (30), and syntenin-1 (SDCBP) (50), but also highlighted differences, such as the relative abundance of some EV markers and the presence of ADP-ribosylation factor 6 (ARF6) and tumor susceptibility gene 101 protein (TSG101) only in MBVs and CM-EVs, respectively (Fig. 4E and Data S3). Hence, our data have identified distinct traits of CM-EVs and MBVs.

To investigate this further, we compared the proteome of pCAF-derived EVs with the proteome of large-medium and small EVs of three publicly available datasets (Fig. S2A-B) (51–53). For each dataset, we selected proteins unique to each EV subpopulation and those with abundance significantly different between the two. Then, we matched this subset to EV proteins whose abundance was significantly different between CM-EVs and MBVs (Data S3). This analysis showed that proteins typically found in large-medium EVs were generally more abundant in MBVs. Proteins typically found in small EVs, instead, were more abundant in CM-EVs (Fig. S2B). This observation was consistent across the three datasets (Fig. S2B). Furthermore, proteins identified only in CM-EVs displayed enrichment for endosome-related GOCC terms (Fig. S2C) that is one of the documented intracellular origins of small EVs (Fig. S2A). Conversely, MBV unique proteins displayed enrichment in GOCC terms associated with plasma membrane, cytosol, ER and mitochondria (Fig. S2C), which are expected in large-medium EVs because of their biogenesis (30). Overall, these results further support that each extracellular compartment contains different subsets of EVs.

The majority of pCAF proteins transferred to ECs during co-culture was identified in both EV types (Fig. 4F, Data S1 and S3), and their abundance positively correlated to the amount measured in the EVs (Fig. 4G, Data S1 and S3). Notably, the majority of the transferred plasma membrane receptors and membrane-bound ligands were more abundant overall in the MBVs (Fig. 4H and Data S3).

Next, we measured whether pCAF-derived CM-EVs and MBVs could transfer proteins to ECs. As these EV types exist in different extracellular sites, we measured protein transfer when CAFs were co-cultured in physical contact (direct co-culture) or not (indirect co-culture) with HUVECs (Fig. 5A). In the direct co-culture, HUVECs are exposed to both types of EVs, while in the indirect co-culture to CM-EVs only (Fig. 5A). Strikingly, the amount of transferred proteins in the direct co-culture was more than two-fold higher compared with the indirect one (Fig. 5B). In line with this result, HUVECs received significantly more proteins when treated with MBVs than with CM-EVs, although the difference was less pronounced (Fig. 5C).

**Fig. 5.**
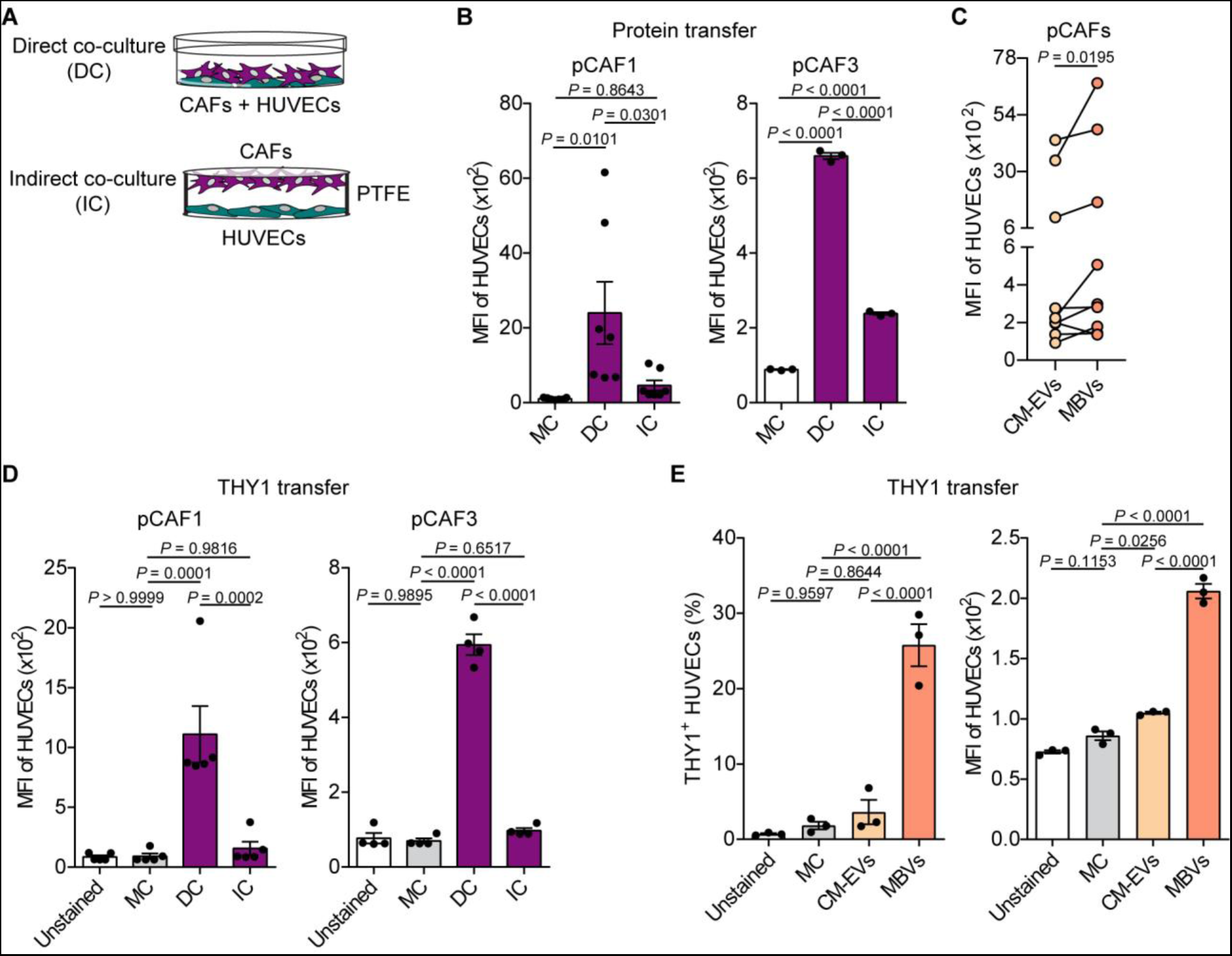
MBVs have a central role in the protein transfer. **(A)** Representation of the co-culturing methods. **(B)** Quantification of the protein transfer from pCAFs to HUVECs in the direct and indirect co-culture, *N =* 7 (pCAF1) or 3 (pCAF3). **(C)** Quantification of the amount of proteins transferred by pCAF-derived CM-EVs and MBVs to HUVECs, *N =* 9. Data are normalized to monoculture. The data related to pCAF-derived MBVs also are shown in Fig. 6B. **(D)** Quantification of THY1 protein levels in monoculture of HUVECs and HUVECs that were directly and indirectly co-cultured with pCAFs, *N =* 5 (pCAF1) and 4 (pCAF3). **(E)** Quantification of THY1^+^ HUVECs and THY1 protein levels in monoculture of HUVECs and in HUVECs treated with pCAF-derived CM-EVs and MBVs, *N =* 3. **(B, D, E)** Data are represented as mean ± SEM, one-way ANOVA with Tukey’s multiple comparison test. **(C)** Two-tailed Wilcoxon matched-pairs test.

Our MS analysis showed that THY1 was significantly more abundant in MBVs than in CM-EVs (Fig. 4H). We used direct and indirect co-cultures to measure THY1 transfer from pCAFs to HUVECs. As for the total proteins, the transfer of THY1 mainly occurred when cells were in direct culture (Fig. 5D). Moreover, upon treatment with MBVs, five-fold more HUVECs positively stained for THY1 compared with when treated with CM-EVs, and THY1 levels were two-fold higher (Fig. 5E). Together, our data provide evidences that MBVs are a major vehicle for protein transfer from mammary CAFs to HUVECs.

### α-SMA^high^ TNFRSF12A^high^ CAFs are the major donors of proteins to ECs

Previous studies have shown that human normal fibroblasts (NFs) activated upon treatment with CM of prostate and melanoma cancer cells transfer more proteins compared with untreated fibroblasts (21). Therefore, we compared protein transfer of our mammary CAFs with their matched NFs isolated from the same patient (pNFs). pNFs are normal-like fibroblasts as they were derived from macroscopically healthy tissue adjacent to the tumor. We found that pCAFs transferred more proteins to HUVECs than pNFs (Fig. 6A), and confirmed this result using microvascular endothelial cells (MVECs) (Fig. S3A). Since we showed that EVs are involved in protein transfer, we compared the amounts of EVs released by pCAFs and pNFs. Nanoparticles Tracking Analysis showed that pCAFs deposited significantly more medium-large EVs in the ECM than their NF counterpart (Fig. 4A, C-D). In the CM, instead, pNFs and pCAFs released EVs of similar size and quantity. (Fig. 4C-D). Notably, HUVECs treated with CAF-derived MBVs received more proteins than when treated with MBVs produced by pNFs (Fig. 6B) and CM-EVs and MBVs secreted by pNFs transferred a comparable amount of proteins to HUVECs (Fig. 6C). Together, these results suggest that CAFs transfer more proteins because they produce more MBVs.

**Fig. 6.**
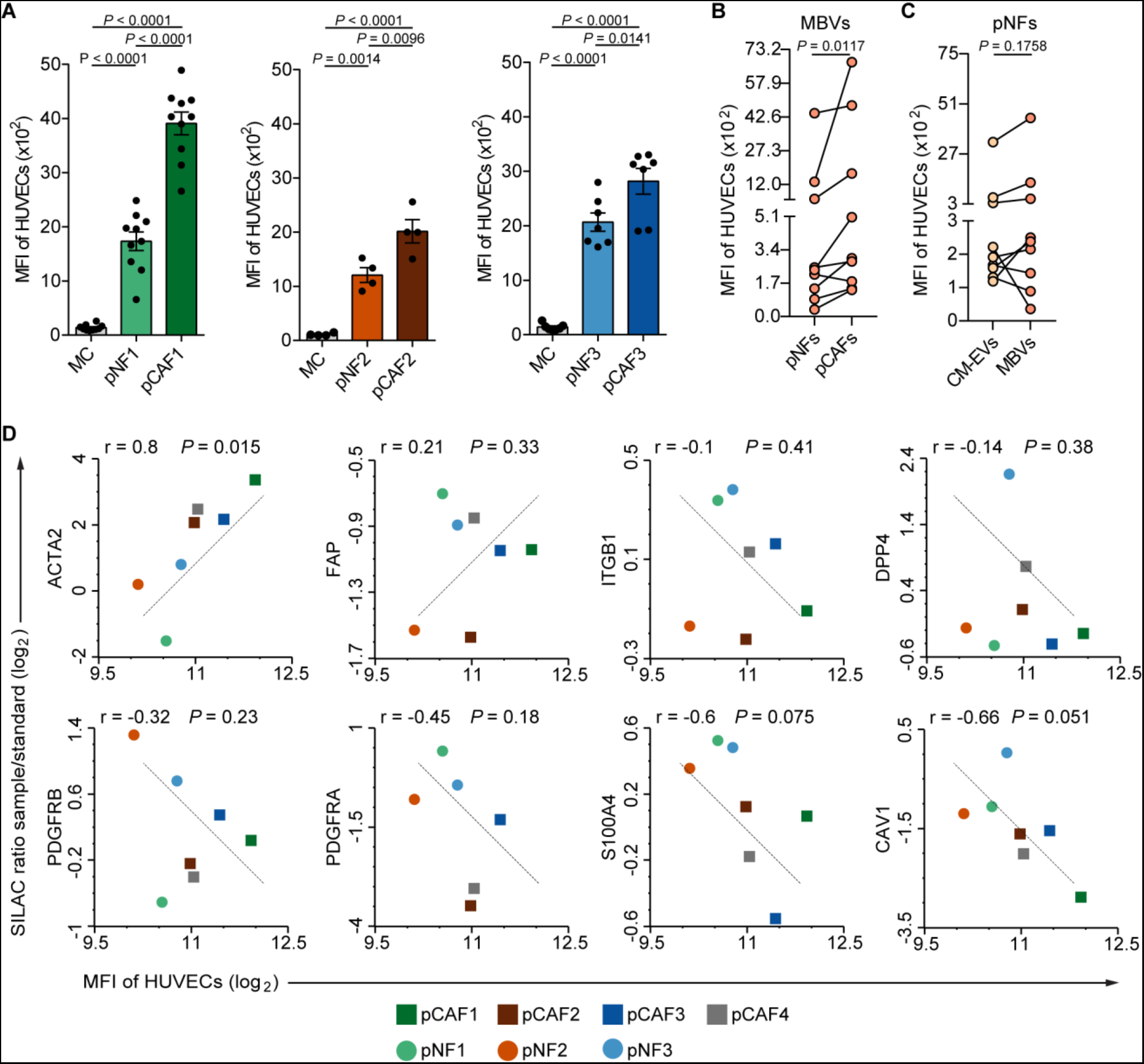
CAFs have an enhanced protein transfer ability. **(A)** Quantification of the protein transfer from pCAFs or pNFs to HUVECs, *N =* 10 (pCAF1 and pNF1), 4 (pCAF2 and pNF2), and 7 (pCAF3 and pNF3). **(B)** Quantification of the amount of proteins transferred by pNF- and pCAF-derived MBVs to HUVECs, *N =* 9. Data are normalized to monoculture. The data related to pCAF-derived MBVs also are shown in Fig. 5C. **(C)** Quantification of the amount of proteins transferred by pNF-derived CM-EVs and MBVs to HUVECs, *N =* 9. Data are normalized to monoculture. The data related to pNF-derived MBVs also are shown in Fig. 6B. **(D)** Scatter plot showing the correlation between the abundance of CAF markers in fibroblasts (Data S4) and the amount of proteins that they transferred to HUVECs, which corresponds to the data shown in Fig. S3B. Data are in the log_2_ scale (r, Pearson correlation). **(A)** Data are represented as mean ± SEM, one-way ANOVA with Tukey’s multiple comparison test. **(B, C)** Two-tailed Wilcoxon matched-pairs test.

Our pCAF lines transferred different amounts of proteins to ECs (Fig. S3B) raising the question of whether all CAFs are able to transfer proteins. To address this question, first we measured the correlation between the abundance of common CAF markers in our pCAF lines (Data S4), including α-SMA, prolyl endopeptidase FAP (FAP), integrin beta-1 (ITGB1), dipeptidyl peptidase 4 (DPP4), platelet-derived growth factor receptor alpha and beta (PDGFRA and PDGFRB), protein S100-A4 (S100A4, also known as FSP-1) and caveolin-1 (CAV1), with their protein transfer ability. Intriguingly, we found that the amount of proteins transferred by fibroblasts significantly correlated (*P* < 0.05) only with α-SMA protein levels (Fig. 6D). Microscopy analysis for α-SMA of our pCAF lines confirmed the proteomic data showing that the pCAF1 line, which transferred more proteins to ECs (Fig. S3B), contained more cells with high α-SMA protein levels than pCAF3 and pCAF4 lines (Fig. 7A-B). To assess whether pCAFs expressing high or low protein levels of α-SMA had different protein transfer ability, we needed to identify cell surface proteins to sort the two living subpopulations for functional assays. To achieve this, we sorted CAFs with high and low α-SMA protein levels and analyzed them by MS proteomics (Fig. S4A). Principal component analysis of 2,080 proteins quantified across the three pCAF lines separated the α-SMA^low^ and α-SMA^high^ subpopulations (Fig. 7C and Data S5). Moreover, 67 proteins showed difference in abundance between α-SMA^low^ and α-SMA^high^ subpopulations in at least two of the three pCAF lines (Fig. S4B and Data S5). Among those, there were 7 cell surface receptors (Fig. 7D). We followed up on the tumor necrosis factor receptor superfamily member 12A (TNFRSF12A, also known as FN14, TweakR or CD266), because its levels were highly different between α-SMA^low^ and α-SMA^high^ CAFs and it was a good candidate for cell sorting (Data S5). Immunofluorescence staining for α-SMA confirmed that there were more α-SMA^high^ cells in TNFRSF12A^high^ sorted pCAFs than in TNFRSF12A^low^ pCAFs (Fig. 7E). Strikingly, TNFRSF12A^high^ pCAFs on average transferred double the amount of proteins to HUVECs than TNFRSF12A^low^ pCAFs (Fig. 7F), including THY1 (Fig. 7G). Notably, TNFRSF12A^low^ pCAFs had a protein transfer ability similar to that of their NF counterpart (Fig. 7F).

**Fig. 7.**
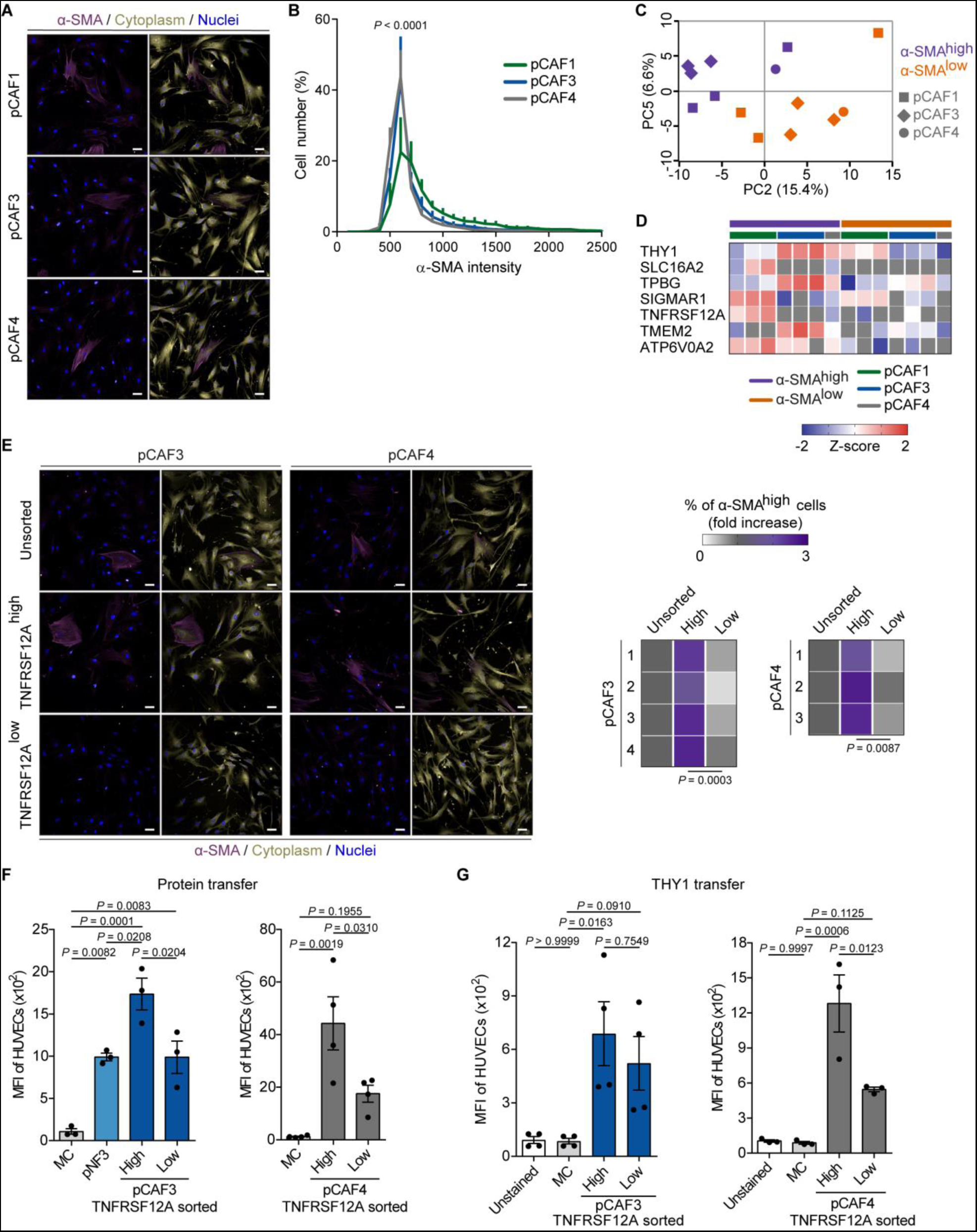
CAFs with high protein transfer ability are a-SMAhigh and TNFRSF12Ahigh. **(A)** presentative images (maximum intensity projection processing from confocal z-stack) of a-SMA ining in pCAFs. Cytoplasm and nuclei were stained with HCS CeiiMask and DAPI, respectively (scale bar = 50 μm). **(B)** Frequency plot obtained from the analysis of the α-SMA staining, it shows the distribution of pCAFs across the different values of α-SMA intensity, *N =* 4. **(C)** Principal component analysis based on 2,080 proteins identified across all the pCAF subpopulations. **(D)** Heatmap based on the Z-score of the LFQ intensity (log_2_) of the cell surface proteins identified in pCAFs (subset of proteins from Fig. S4B), *N =* 7. **(E)** Representative images (maximum intensity projection processing from confocal z-stack) of α-SMA staining in unsorted, TNFRSF12A^high^ and TNFRSF12A^low^ pCAFs. Cytoplasm and nuclei were stained with HCS CellMask and DAPI, respectively (scale bar = 50 μm). The heatmap shows the increase in the percentage of α-SMA^high^ pCAFs in sorted pCAFs with respect to the unsorted ones. **(F)** Quantification of the protein transfer from TNFRSF12A^high^ pCAFs, TNFRSF12A^low^ pCAFs or pNFs to HUVECs, *N =* 3 (pCAF3 and pNF3) or 4 (pCAF4). **(G)** Quantification of THY1 protein levels in monoculture of HUVECs and HUVECs that were co-cultured with TNFRSF12A^high^ or TNFRSF12A^low^ pCAFs, *N =* 4 (pCAF3) or 3 (pCAF4). **(B)** Two-way ANOVA test with Tukey’s multiple comparison test, it compares pCAF3 or pCAF4 with pCAF1 at α-SMA intensity values of 500 and 600. **(E)** Unpaired t test with Welch’s correction. **(F, G)** Data are represented as mean ± SEM, one-way ANOVA with Tukey’s multiple comparison test.

Hence, α-SMA^high^ mammary CAFs can be enriched using the transmembrane receptor TNFRSF12A and have enhanced ability to transfer proteins to ECs.

### α-SMA^high^ TNFRSF12A^high^ CAFs express high levels of myofibroblast markers

α-SMA^high^ CAFs are typically those referred to as myCAFs, while α-SMA^low^ are typically the iCAFs (15, 16). Therefore, we investigated the expression of other myCAF and iCAF markers in our CAF subpopulations. To do this, we sorted TNFRSF12A^high^ and TNFRSF12A^low^ pCAFs, expanded them in culture, and then assessed the expression of CAF markers by RT-qPCR. This analysis confirmed that TNFRSF12A^high^ pCAFs expressed higher levels of *ACTA2* and other genes highly expressed in mammary myCAFs (7, 16, 54), such as collagen alpha-1 (I) chain (*COL1A1*) and transgelin (*TAGLN*), compared with TNFRSF12A^low^ pCAFs (Fig. S5A-B). Conversely, we did not detect significant differences in mRNA levels of stromal cell-derived factor 1 (*SDF1*, also known as C-X-C motif chemokine 12 or *CXCL12*) and interleukin-6 (*IL6*), which are highly expressed in mammary iCAFs (7, 16, 54) (Fig. S5A-B). Similarly, decorin (*DCN*), which has been found expressed at similar levels in all CAFs (7, 54), had similar mRNA levels in our two sorted populations (Fig. S5A-B). These data suggest that high levels of the TNFRSF12A receptor are found in CAFs with myCAF phenotype. Consistent with this observation, in two publicly available single-cell RNA sequencing datasets of CAFs isolated from patients with BC (7, 16), we found that both *ACTA2* and *TNFRSF12A* mRNA levels were high in the subpopulation defined by the authors as myCAFs (Fig. 8A-B). In addition, immunofluorescence staining of tumor tissue sections from patients with BC confirmed the presence in the stroma of TNFRSF12A^+^ and α-SMA^+^ CAFs and showed that these cells can exist in close proximity to blood vessels (Fig. 8C and Fig. S6). Hence, enhanced CAF-EC communication based on protein transfer is distinctive of those CAFs with a myofibroblast-like phenotype.

**Fig. 8.**
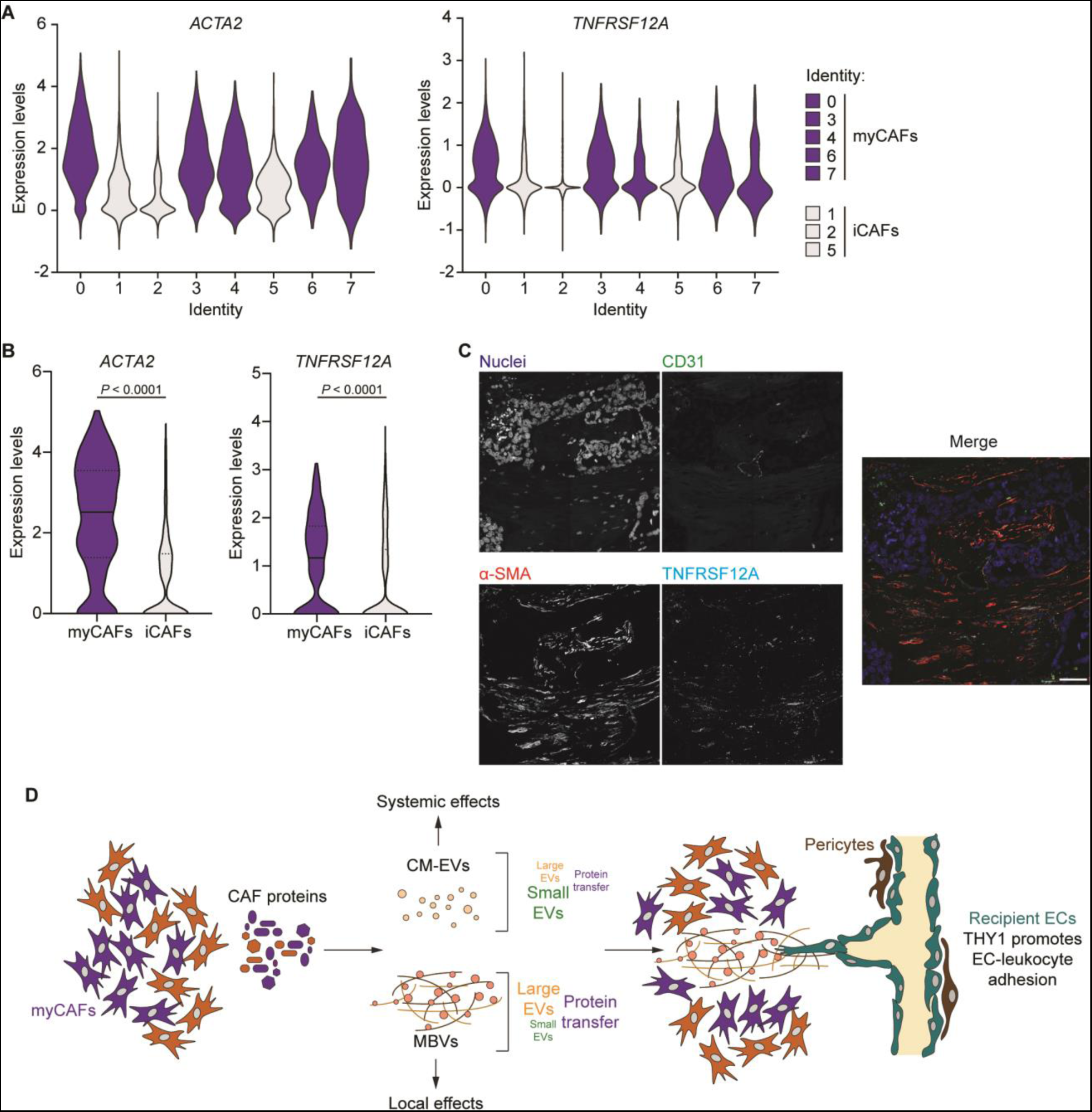
Characterization of CAFs with high protein transfer ability. **(A, B)** Violin plot showing the expression levels of *ACTA2* and *TNFRSF12A* in the myCAF and iCAF subpopulations, **(A)** data are from (16), **(B)** data are from (7). **(C)** Representative image of TNFRSF12A, α-SMA and CD31 staining in a tumor tissue section from a patient with BC (maximum Z projection). Nuclei were stained with Hoechst-33342, scale bar = 50 μm. **(D)** Working model showing the myCAF-EC communication based on the MBV-mediated transfer of proteins.

## Discussion

Using CAFs isolated from patients with breast cancer, we have discovered that CAFs with a myofibroblastic-like phenotype transfer high amounts of proteins to the surrounding endothelium. The transfer of proteins mainly occurs through matrix-bound vesicles. Using THY1 as an example of transferred protein, our work also shows that transferred proteins can influence the phenotype of the endothelium (Fig. 8D). Many works have shown that CAFs in different states secrete distinct subsets and amounts of soluble factors and ECM components, which determine their functions in the tumor (15). Therefore, understanding the heterogeneity of CAFs by associating their states to specific biological functions is fundamental to design drugs for cancer treatment.

It is known that CAFs use EVs (14) and intercellular transfer of proteins to affect the function of neighboring cells in vitro (21, 22). While these mechanisms have been originally described between CAFs and cancer cells, using various MS-based proteomic approaches we show that ECs also receive proteins from CAFs in vitro. Moreover, we provide evidences that this process may occur in vivo. What is the fate of these proteins in recipient cells? It has been suggested that the fate of the EV cargo depends on the mechanism of EV internalization (55, 56). For example, the EV cargo can be directed toward lysosomes for degradation or can escape it (55–57). Here, we show that CAF-derived proteins can be functional in recipient cells: ECs can receive plasma membrane proteins from CAFs, most of which are involved in migration and cell-cell or cell-matrix adhesion. Among them, THY1 enhanced the ability of ECs to interact physically with THP-1 monocytes. This result is in line with other studies showing that proteins transferred by cancer cells or CAFs influence the phenotype of recipient cells (21, 22, 47–49). Interestingly, it has been previously shown that PC3 human prostate cancer cells increase the migration of prostate cancer and benign prostatic hyperplasia cells via exosomal transfer of integrin αvβ3 (47). Moreover, HeyA8 and TYK-nu human epithelial ovarian cancer cells induce an invasive mesenchymal phenotype of human peritoneal mesothelial cells through the transfer of exosomal CD44 (48), and activated human prostate and dermal fibroblasts support migration of cancer cells by transferring galectin-1 through ectosomes (22). Notably, we found that CAFs transferred all these proteins to ECs, and that CD44 and ITGB3 were among the receptors with the highest exogenous fraction. This suggests that, in addition to THY1, CD44 and ITGB3 may also influence EC functions.

The ECM is an important source of signals that actively regulate tumor progression. Its structure and composition, including ECM-associated proteins such as growth factors, control most if not all aspects of tumor pathology (58–60). It is now evident that EVs have an essential role in the ECM biology: EVs can be functional components of the ECM (32, 34) and CM-EVs control ECM deposition (61) and remodeling (62). Our work provides a first evidence that CAFs deposit EVs in the matrix and that MBVs play a key role as vehicles for intercellular protein transfer. MBVs and CM-EVs contain EVs of different sizes, and our MS proteomic characterization identified several our work supports the concept that CM-EVs and MBVs may have different intracellular origins, specifically endosomal for CM-EVs and plasma membrane for MBVs, and provides initial evidence that these two subsets of vesicles have distinct functions. We show that CAFs secrete more MBVs than their normal-like fibroblast counterpart. This may explain why CAFs have a greater capacity to transfer proteins to the endothelium. Based on this, we argue that protein transfer from fibroblasts is a phenomenon predominant in pathological conditions. Our work also indicates that CAF-derived MBVs may play unique roles in altering tumors locally, in addition to being a source of systemic signals, which is the more classical function associated with tumor-derived EVs (63). myCAFs deposit the majority of the tumor ECM and contribute to its remodeling (16, 60). Our work suggests that there are additional ways through which myCAFs can alter the tumor ECM and influence cellular function. We show that myCAFs are the major donors of proteins to ECs, but we cannot exclude that it depends on their ability to deposit in the matrix different types and/or amounts of MBVs, or on their ability to remodel the ECM and to make MBVs more available for uptake by ECs, or on their ability to produce large quantities of ECM proteins that can bind a bigger amount of EVs to the matrix. Another interesting aspect that emerged from our work is that MBVs could be a source of nutrients for tumor and stromal cells. It is known that cancer cells (20, 64, 65) and ECs (66) take up proteins and amino acids from the extracellular milieu and use them for macromolecule synthesis, and to modulate redox homeostasis (67). In recipient cells, CAF-derived proteins may undergo proteolytic degradation and supply amino acids that contribute to biosynthetic and bioenergetics processes. Hence, we would speculate that MBVs can be an additional mechanism for local exchange of nutrients within the TME (68).

Finally, we have identified TNFRSF12A as potential cell surface marker of mammary CAFs with myofibroblast-like phenotype to be added to the small panel of plasma membrane proteins that can be used to isolate these cells for functional characterization (15). Indeed, after the exclusion of epithelial cells, immune cells, ECs and pericytes, the TNFRSF12A receptor allows the direct selection of CAFs expressing high levels of myofibroblast markers.

In conclusion, our work has identified a novel function of myCAFs: their ability to transfer functional proteins to ECs through MBVs. Our work paves the way for studies aimed at exploring whether and how other transferred plasma membrane proteins can modify endothelial cell phenotype, and how this affects the function of the tumor vasculature and influences tumor development and progression in vivo. This will inform us on whether targeting the production of MBVs should be further investigated as a strategy to oppose cancer. For that, more work should also be done to understand MBV biogenesis and function, a mechanism that so far has been largely overlooked.

## Limitations

To test the transfer of proteins in vivo we have used mice that express RFP under the control of the α-SMA promoter, and we cannot exclude that α-SMA expressing cells other than CAFs, for example pericytes surrounding the endothelium, also transfer proteins to ECs (69, 70). However, CAFs expand during tumor progression and we have shown that the amount of transferred proteins to ECs increases proportionally with CAF number, in vitro. Hence, we think that the majority of RFP found in the tumor endothelium could derive from CAFs. Another limitation is that we cannot exclude that CAFs had transferred RFP mRNA, which has then been translated into protein in the endothelium. Hence, additional studies are needed to further prove that CAFs transfer proteins to the endothelium in vivo, for example using MetRS* mice (71).

## Materials and Methods

### Cell culture

pCAFs and pNFs were isolated at the CRUK Beatson Institute from women with breast cancer and immortalized as previously described (35). Samples were obtained through NHS Greater Glasgow and Clyde Bio-repository. Patients agreed with the use of their tissue samples for research. pCAFs and pNFs were cultured on dishes coated with collagen I from rat tail (12 μg/ml, Gibco, Thermo Fisher Scientific, Waltham, MA). Unless otherwise stated, pCAFs and pNFs used for the experiments are pCAF1 and pNF1. cCAFs and cNFs were kindly provided by Professor Akira Orimo (Juntendo University, Tokyo) (35, 72). CAFs, NFs and luciferase-expressing 4T1 mouse breast cancer line were cultured in Dulbecco’s modified Eagle’s medium (DMEM) supplemented with 10% fetal bovine serum (FBS), 2 mM glutamine and 1% penicillin/streptomycin (Life Technologies, Thermo Fisher Scientific). HUVECs were isolated from donors using previously described methods (59). HMVECs (100-05a) were purchased from Sigma-Aldrich (St. Louis, MO). HUVECs and HMVECs were cultured on 1% gelatin coated dishes in EGM-2 or EGM-2 MV (Lonza, Basel, Switzerland), respectively. THP-1 cells were purchased from ATCC and cultured in RPMI-1640 medium (Life Technologies) supplemented with 10% FBS, 2 mM glutamine and 0.05 mM 2-mercaptoethanol (Sigma-Aldrich). MCF10DCIS.com cells (35) were cultured in F12 medium supplemented with 5% horse serum (Life Technologies), 2 mM glutamine and 1% penicillin/streptomycin. Jurkat cells were cultured in RPMI-1640 medium supplemented with 10% FBS and 2 mM glutamine. For SILAC experiments, fibroblasts were cultured in SILAC DMEM (Life Technologies) supplemented with 2% FBS, 8% 10 kDa dialyzed FBS, 2 mM glutamine, 1% penicillin/streptomycin, 84 mg/l ^13^C_6_^15^N_4_ L-arginine and 175 mg/l ^13^C_6_^15^N_2_ L-lysine (Cambridge Isotope Laboratories, Inc., Tewksbury, MA). All the cell lines were cultured under standard conditions (37°C and 5% CO_2_) and were routinely tested for mycoplasma.

### EV isolation

pCAFs or pNFs were plated in serum-free DMEM. After 48h, the EVs were collected from both the CM and the matrix. The EVs from the matrix were detached using Accutase^®^ (Sigma-Aldrich) and collected in DMEM supplemented with 0.5% FBS, which had been previously ultracentrifuged at 100,000 x g for 5h to reduce the amount of serum EVs. Then, cells and debris were removed by centrifugation at 300 x g (4°C, 10min) and 2,000 x g (4°C, 30min), and CM-EVs and MBVs were isolated by ultracentrifugation at 100,000 x g (4°C, 90min). Pelleted EVs were resuspended in PBS and subjected to another step of ultracentrifugation at 100,000 x g (4°C, 90min). Then, EVs were collected in PBS and gently sonicated at 5 microns amplitude using a metal tip (Soniprep 150, MSE) three times for 5s before using them for protein transfer experiments and nanoparticles tracking analysis and before sample preparation for electron microscopy.

### Small interfering RNA

Transient knockdown was performed using the Amaxa kit R (Lonza) and Nucleofector device (program T-20, Lonza) according to the manufacturer’s protocol. 2×10^6^ CAFs were transfected with 3 nM of non-targeting control (D-001810-10-05, GE Healthcare Dharmacon, Inc., Lafayette, CO) or THY1 human siRNA (L-015337-00-0005, GE Healthcare Dharmacon, Inc.). CAFs were used for experiments 72h after transfection.

### Nanoparticles tracking analysis

CM-EVs and MBVs were isolated from 1.5×10^5^ pCAFs or pNFs, which were seeded in serum-free medium in 6 cm cell culture dishes and isolated as described above. After isolation, the EVs were resuspended in 1 ml of PBS. EV size and concentration were determined using a NanoSight LM10 (Malvern Panalytical, Malvern, UK) and the NTA 3.1 software. Each measurement is the result of three acquisitions of 60s. The camera level was set to 14 and the detection threshold to 4. The PBS used for EV isolation and collection was filtered with the 0.02 μm filter.

### Intercellular protein and THY1 transfer

To measure the intercellular protein transfer in co-culture, donor cells were labeled with 10 μM CellTrace^TM^ CFSE (Life Technologies) in PBS for 20min at 37°C. After at least 1h, donor cells were seeded. Once they were adhered, recipient cells were seeded in monoculture and co-culture with donor cells for 24h. In the indirect co-culture, pCAFs (donor cells) were plated on glass coverslip, which was positioned upside down on the culture dish where HUVECs were seeded. The glass coverslip was placed above a polytetrafluoroethylene film (0.08 mm thickness, Goodfellow Cambridge Ltd, Huntingdon, UK). After 24h of co-culture, cells were detached with Accutase^®^, resuspended in FACS buffer (25 mM HEPES, 5 mM EDTA, 1% penicillin/streptomycin, 1% FBS in PBS) and the transfer of CFSE labeled proteins was analyzed by an Attune^TM^ NxT flow cytometer (Thermo Fisher Scientific) and FlowJo software version 10.7.1. To measure THY1 transfer, cells were detached with Accutase^®^ after 24h of co-culture, resuspended in FACS buffer, incubated with APC anti-THY1 antibody (1/100, [5E10] 328114 BioLegend, San Diego, CA) and human TruStain FcX^TM^ (1/200, BioLegend) for 45min on ice (100 μl/10^6^ cells). DAPI (Sigma-Aldrich) was used as live/dead marker. THY1 transfer was analyzed by an Attune^TM^ NxT flow cytometer and FlowJo software version 10.7.1. Unless otherwise stated, donors and recipient cells were plated at 2:1 ratio. The medium used for the co-culture experiments was EGM-2 or EGM-2 MV depending on the EC type.

To measure the EV-mediated transfer of proteins, CM-EVs and MBVs were isolated from 1.5×10^5^ pCAFs or pNFs and labeled with 10 μM CellTrace^TM^ CFSE in PBS for 20min at 37°C. After labeling, EVs were washed in PBS by two sequential steps of ultracentrifugation at 100,000 x g (4°C, 90min). The isolated EVs were used to treat 3×10^4^ HUVECs in EGM-2. After 20h, HUVECs were detached and intercellular protein transfer was analyzed as above. To measure the EV-mediated transfer of THY1, CM-EVs and MBVs were isolated from 5×10^4^ pCAFs and used to treat 2.5×10^4^ HUVECs in EGM-2. After 3h, HUVECs were detached, stained for THY1 and analyzed as above.

### Western blotting analysis

Cells were lysed in 2% SDS in 100 mM Tris-HCl pH 7.4, incubated at 95°C for 5min, sonicated using a metal tip and centrifuged at 16,000 x g for 10min. Protein concentration was determined using Optiblot Bradford reagent (Abcam, Cambridge, UK). Protein lysate was mixed with NuPAGE^TM^ LDS sample Buffer (4x) (Life Technologies) supplemented with 400 mM dithiothreitol (DTT, Sigma-Aldrich). Proteins (15-20 μg) were separated using 4-12% gradient NuPAGE^TM^ Novex Bis-Tris gel (Life Technologies). Protein transfer was performed on methanol-activated Immobilon^®^-FL PVDF membrane (Sigma-Aldrich). Membrane was blocked for 1h in 5% BSA (bovine serum albumin, Sigma-Aldrich) in Tris-buffered saline with 0.1% Tween^®^ 20 detergent (TBST) at room temperature and incubated with the primary antibody overnight at 4°C. The following primary antibodies were used: THY1 (1/2,000, [D3V8A] 13801 Cell Signaling Technology, Danvers, MA) and GAPDH (1/1,000, sc-48167 Santa Cruz Biotechnology, Inc., Dallas, TX). The membrane was incubated with HRP-conjugated (1/2,500 New England Biolabs, Ipswich, MA) or IRDye^®^ (1/10,000 LI-COR Biosciences, Lincoln, NE) antibody for 45min at room temperature. Western blot images were acquired using a myECL Imager (Thermo Fisher Scientific) or a LI-COR Odyssey CLx scanner (Image Studio software, version 5.0.21).

### Immunofluorescence

For vimentin staining, cultured pCAFs and pNFs were fixed in 4% paraformaldehyde (PFA, Sigma-Aldrich), permeabilized with 0.1% Triton X-100 (TX-100, Sigma-Aldrich) in 1% BSA in PBS for 30min, incubated overnight at 4°C with anti-vimentin antibody (1/50, sc-7557 Santa Cruz Biotechnologies) and then with Alexa Fluor^®^ 488 secondary antibody (1/400, Life Technologies) for 2h. DAPI (1/5,000) was used for nuclear staining. Images were acquired using a Zeiss LSM 710 confocal microscope (Carl Zeiss, Oberkochen, Germany, EC Plan-Neofluar 20x/0.50 M27 objective, no immersion).

To evaluate protein transfer by immunofluorescence, pCAFs were labeled with CellTrace^TM^ CFSE as described above and seeded on glass coverslips in a 24-well plate. Once they were adhered, ECs were seeded in monoculture or in co-culture with pCAFs. After 24h, cells were fixed in 4% PFA. DAPI and Alexa^®^ 647 Phalloidin (1/100, Life Technologies) were used for nuclear and F-actin staining, respectively. Images were acquired with a Zeiss LSM 880 confocal microscope in Airyscan mode (Carl Zeiss, Plan-Apochromat 63x/1.4 Oil DIC M27 objective, zoom 1.8, z-stacks of 5-9 μm, 28-48 slices). Images were Airyscan processed with Zen software using default settings. The 3D reconstruction and analysis were performed using Imaris software (version 9.5, Bitplane, Oxford Instruments, Abingdon, UK).

For α-SMA staining, 5×10^3^ pCAFs were seeded in each well of a 96-well plate. The following day, cells were fixed in 50% acetone/50% ethanol for 20min and permeabilized with 0.05% saponin (Sigma-Aldrich) in 1% BSA in PBS for 30min. Cells were incubated with anti-α-SMA antibody (1/200, [1A4] ab7817 Abcam) in 1% BSA in PBS for 1.5h and then with Alexa Fluor^®^ 647 secondary antibody (1/250, Life Technologies) for 1h. HCS CellMask^TM^ Green Stain (1/10,000, Life Technologies) and DAPI were used for cytoplasm and nuclear staining, respectively. For each well, 45 images were acquired on an Opera Phenix high-content imaging system (20x objective, z-stacks of 0.8 μm, PerkinElmer, Waltham, MA). Image analysis was performed using Harmony imaging analysis software (PerkinElmer, version 4.9).

### pCAF sorting

Cultured pCAFs were detached with Accumax^TM^ solution (Sigma-Aldrich) and resuspended in FACS buffer.

pCAF sorting based on α-SMA protein level: pCAFs were fixed and permeabilized by using the eBioscience^TM^ Intracellular Fixation & Permeabilization Buffer Set (Life Technologies) according to the manufacturer’s instructions. Cells were resuspended in permeabilization buffer (100 μl/10^6^ cells) and incubated with anti-α-SMA antibody (1/1,000, [1A4] ab7817 Abcam) for 1h and then with Alexa Fluor^®^ 488 or 647 secondary antibody (1/250, Life Technologies) supplemented with 2% donkey serum for 1h.

pCAF sorting based on TNFRSF12A protein level: 10^6^ cells were incubated with BD Horizon BV421 anti-TNFRSF12A antibody (1/140, 565712 BD Biosciences, Franklin Lakes, NJ) and human TruStain FcX^TM^ (1/200, BioLegend) in 100 μl of FACS buffer for 45min on ice. pCAFs were sorted in α-SMA^high^ and α-SMA^low^ or TNFRSF12A^high^ (top 10%) and TNFRSF12A^low^ (bottom 10%) using a BD FACSAria^TM^ (BD Biosciences).

### Adhesion Assay

Control and THY1-silenced pCAFs were labeled with 2.5 μM CellTracker^TM^ Green CMFDA Dye (Life Technologies) in PBS for 25min at 37°C and 2×10^4^ labeled pCAFs were seeded in a gelatin coated well of a 96-well plate. HUVECs were labeled with 1 μM CellTracker^TM^ Deep Red Dye (Life Technologies) in PBS for 20min at 37°C and 4×10^4^ labeled cells were seeded in co-culture with pCAFs. After 24h of co-culture, 8.5×10^3^ THP-1 cells, which were labeled with 2 μM CellTracker^TM^ Orange CMTMR Dye (Life Technologies) in PBS for 20min at 37°C, were added in each well in M199 medium (Life Technologies) supplemented with 10% FBS. After 45min, unbound THP-1 cells were removed by three washes in 1% BSA in PBS with calcium and magnesium (Sigma-Aldrich). Cells were fixed in 4% PFA and DAPI was used for nuclear staining. For each well, 25-45 images were acquired on an Opera Phenix high-content imaging system (objective 20x, z-stacks of 2 μm, PerkinElmer). Image analysis was performed using Harmony imaging analysis software (PerkinElmer, version 4.9). Only the THP-1 cells overlapping the ECs at least for the 30% of their cellular body were counted.

### CAF characterization

pCAFs, MCF10DCIS.com cells and HUVECs were detached with the Accumax^TM^ solution and resuspended in FACS buffer. Jurkat cells were centrifuged and resuspended in FACS buffer. 10^6^ cells were incubated with PE anti-CD31 ([WM59] 303105), PE anti-CD45 ([2D1] 368510) and PE anti-CD326 (EpCAM, [9C4] 324206) antibodies (1/100, BioLegend) and human TruStain FcX^TM^ (1/200, BioLegend) in 100 μl of FACS buffer for 45min on ice. DAPI was used as live/dead marker. Cells were analyzed using a BD LSRFortessa^TM^ Cell Analyzer (BD Biosciences) and FlowJo software version 10.7.1.

### MS proteomic analysis

For trans-SILAC experiments, heavy-labeled pCAFs were labeled with CellTracker^TM^ Green CMFDA Dye as described above and 1.5×10^6^ pCAFs were seeded in a gelatin coated 15 cm dish. After 16h, 7.5×10^5^ HUVECs were seeded in monoculture and in co-culture with pCAFs. After 4h and 24h, cells were detached with Accutase^®^, resuspended in FACS buffer, sorted using a BD FACSAria^TM^ and lysed in 6 M urea/ 2 M thiourea supplemented with 10 mM tris(2-carboxyethyl)phosphine (TCEP) and 40 mM chloroacetamide (CAA) in 75 mM NaCl and 50 mM Tris-HCl (Sigma-Aldrich) and sonicated using a metal tip. 25-120 μg of proteins were digested with trypsin and were fractionated using high pH reverse phase fractionation. Briefly, dried peptides were resuspended in 200 mM ammonium formate adjusted to pH 10 with ammonium hydroxide solution (Sigma-Aldrich). Then, peptides were loaded on pipette-tip columns of ReproSil-Pur 120 C18-AQ 5 μm (Dr. Maisch HPLC GmbH, Ammerbuch, Germany), eluted in 7 fractions using an increasing amount of acetonitrile and analyzed by MS.

For the proteome analysis of THP-1, α-SMA^high^ and α-SMA^low^ pCAFs, cells were washed three times in PBS and lysed in 6 M urea/ 2 M thiourea supplemented with 10 mM TCEP and 40 mM CAA in 75 mM NaCl and 50 mM Tris-HCl and sonicated using a metal tip. 10 μg of proteins were digested with trypsin, desalted using C18 StageTip (73) and analyzed by MS.

For the analysis of pCAF1 proteome, cultured pCAFs were washed three times with PBS and cultured in serum-free DMEM. After 24h, cells were washed with PBS, lysed in 2% SDS with 1 mM DTT in 100 mM Tris-HCl pH 7.4, incubated at 95°C for 5min and sonicated using a metal tip. Tryptic peptides were generated from 150 μg of proteins using filter-aided sample preparation using filtration units with MW cutoff of 30 kDa (74, 75). Briefly, lysates were loaded on the filter units, incubated for 20min with 55 mM iodoacetamide (IAA, Sigma-Aldrich) in 50 mM ammonium bicarbonate pH 8.0, digested with trypsin and eluted with 50 mM ammonium bicarbonate pH 8.0 (Sigma-Aldrich). Then, 60 μg of peptides were fractionated using high pH reverse phase fractionation as described above and analyzed by MS.

For the analysis of the EV proteome, pCAFs were cultured in DMEM supplemented with 10% ultracentrifuged FBS. Then, pCAFs were washed three times in PBS and cultured in serum-free DMEM. After 48h, CM-EVs and MBVs were collected as described above from 2×10^7^ and 10^7^ pCAFs, respectively. After ultracentrifugation, EVs were collected in 200 mM HEPES pH 8.0 and 2,2,2-Trifluoroethanol (TFE) was added 1:1 (Sigma-Aldrich). EVs were sonicated three times at 10 microns amplitude for 10 seconds (with 20 seconds on ice in between each sonication), then EVs were incubated at 60°C for 1h at 1000 rpm and sonicated again. EV lysates were incubated with 10 mM TCEP and 40 mM CAA for 1h. Then, TFE concentration was reduced to 10% by adding 200 mM HEPES and EV lysates were digested with trypsin. Peptides were desalted using C18 StageTip, dried, resuspended in 200 mM HEPES and incubated with 0.1 mg Tandem Mass Tag (TMTzero^TM^, Thermo Fisher Scientific) label reagent for 2h at 26°C and 450 rpm. Samples were dried, resuspended in 0.1% formic acid, acidified by adding trifluoroacetic acid (TFA, Sigma-Aldrich), desalted using C18 StageTip and analyzed by MS.

For the proteome analysis of pCAFs and pNFs, cultured cells were washed with PBS, lysed in 2% SDS in 100 mM Tris-HCl pH 7.4, incubated at 95°C for 5min, sonicated using a metal tip and centrifuged at 16,000 x g for 10min. Lysates were mixed 1:1 with an internal standard composed of a mix of SILAC heavy-labeled cCAFs/cNFs. Protein lysate was mixed with NuPAGE^TM^ LDS sample Buffer (4x) and 1 mM DTT. Proteins were separated using 4-12% gradient NuPAGE^TM^ Novex Bis-Tris gel, which then was stained with Coomassie Blue. Gel lanes were cut into slices, proteins were in gel digested with trypsin (76), peptides were desalted using C18 StageTip and analyzed by MS.

### MS analysis: Q Exactive HF (trans-SILAC experiment and pCAF1 proteome)

Each of the 7 fractions was dried down and then re-suspended in 2% acetonitrile/0.1% TFA in water and separated by nanoscale C18 reverse-phase liquid chromatography performed on an EASY-nLC II 1200 coupled to a Q Exactive HF mass spectrometer (Thermo Fisher Scientific). Elution was carried out for a total run time duration of 65min (fraction 1), 105min (from fraction 2 to 5) and 135min (fraction 6 and 7), using an optimized gradient. Peptides were subsequently eluted into a 50 cm (trans-SILAC) or 20 cm (pCAF1 proteome) fused silica emitter (New Objective, Inc., Littleton, MA) packed in-house with ReproSil-Pur C18-AQ, 1.9μm resin (Dr. Maisch HPLC GmbH). The emitter was kept at 50°C (trans-SILAC) or 35°C (pCAF1 proteome) by means of a column oven integrated into the nanoelectrospray ion source (Sonation).

Eluting peptides were electrosprayed into the mass spectrometer using a nanoelectrospray ion source (Thermo Fisher Scientific). An Active Background Ion Reduction Device (ABIRD, ESI source solutions) was used to decrease air contaminants signal level.

The Xcalibur software (Thermo Fisher Scientific) was used for data acquisition. Full scans over mass range of 375–1500 m/z were acquired at 60,000 resolution at 200 m/z. Multiply charged ions from two to five were selected through a 1.4 m/z window and fragmented. Higher energy collisional dissociation fragmentation was performed on the 15 most intense ions, using normalized collision energy of 27, and resulting fragments were analyzed in the Orbitrap at 15,000 resolution, using a maximum injection time of 25ms or a target value of 10^5^ ions. Former target ions selected for MS/MS were dynamically excluded for 20s.

### MS analysis: Orbitrap Fusion^TM^ Lumos^TM^ (THP-1, sorted pCAF and EV proteome)

Desalted peptides were separated by nanoscale C18 reverse-phase liquid chromatography performed on an EASY-nLC 1200 coupled to an Orbitrap Fusion^TM^ Lumos^TM^ mass spectrometer (Thermo Fisher Scientific). Elution was carried out for a total run time duration of 265min (THP-1 and sorted pCAF proteome) or 135min (EV proteome), using a binary gradient with buffer A (water) and B (80% acetonitrile), both containing 0.1% of formic acid. Peptide mixtures were separated at 300 nl/min flow, using a 50 cm fused silica emitter (New Objective, Inc.) packed in-house with ReproSil-Pur C18-AQ, 1.9μm resin (Dr Maisch GmbH). Packed emitter was kept at 50°C by means of a column oven integrated into the nanoelectrospray ion source (Sonation). The eluting peptide solutions were electrosprayed into the mass spectrometer via a nanoelectrospray ion source (Sonation). An ABIRD (ESI source solutions) was used to decrease ambient contaminant signal level.

Samples were acquired on an Orbitrap Fusion^TM^ Lumos^TM^ mass spectrometer. The mass spectrometer was operated in positive ion mode and used in data-dependent acquisition mode (DDA). Advanced Peak Determination was turned on and Monoisotopic Precursor Selection was set to “Peptide” mode. A full scan was acquired at a resolution of 120,000 (THP-1 and sorted pCAF proteome) or 60,000 (EV proteome) at 200 m/z, over mass range of 375-1500 m/z (THP-1 and sorted pCAF proteome) or 375-1400 m/z (EV proteome). The top 20 (THP-1 and sorted pCAF proteome) or 15 (EV proteome) most intense ions were selected using the quadrupole, fragmented in the ion routing multipole, and finally analyzed in the linear ion trap (THP-1 and sorted pCAF proteome) or analyzed in the Orbitrap at 15,000 resolution (EV proteome), using a maximum injection time of 35ms (THP-1 and sorted pCAF proteome) or 125ms (EV proteome), or a target value of 2×10^4^ ions (THP-1 and sorted pCAF proteome) or 1.5×10^5^ ions (EV proteome). Former target ions selected for MS/MS were dynamically excluded for 60s (THP-1 and sorted pCAF proteome) or 30s (EV proteome).

### MS proteomic data analysis

The .RAW files were processed with MaxQuant software (version 1.5.5.1 for the proteome analysis of pCAF/pNF proteome, version 1.6.3.3 for all the other experiments) (77) and searched with the Andromeda search engine. The following setting was used: minimal peptide length 7 amino acids, trypsin specific digestion mode with maximum 2 missed cleavages, carbamidomethyl (C) as fixed modification, and oxidation (M) and acetylation (Protein N-term) as variable modifications. For the analysis of EV proteome, TMTzero^TM^ was added as fixed modification and maximum 4 missed cleavages were allowed. Minimum peptide ratio count was set to 2, except that for the trans-SILAC experiment and the analysis of pCAF1 proteome where this parameter was set to 1. “Unique + razor” peptides were used for quantification in the analysis of THP-1 and pCAF/pNF proteome; “unique” peptides were used for quantification in all the other experiments. The “match between runs” option was enabled for the analysis of pCAF/pNF proteome. For SILAC experiments, multiplicity was set to 2: light labels were Arg0 and Lys0; heavy labels were Arg10 and Lys8. Label free quantification (LFQ) setting was enabled for all the other experiments. The false discovery rates (FDRs) at protein and peptide levels were set to 1%.

Perseus software (version 1.5.5.3 for the analysis of pCAF/pNF proteome and version 1.6.2.2 for all the other experiments) (78) was used for data analysis. Potential contaminants, reverse peptides and proteins only identified by a modification site were filtered out. Only proteins identified with at least one unique peptide were kept for the analysis. To define the transferred proteins in the trans-SILAC experiment, we selected proteins that have a “Ratio H/L count” value higher in HUVECs co-cultured with CAFs compared with monoculture. In addition, we selected proteins that have an intensity value in the heavy channel (Intensity H) but not in the light one (Intensity L). The exogenous fraction was calculated as: 1-[1/(x+1)], where x is the “Ratio H/L” value. The exogenous fraction of proteins that have an “Intensity H” value but not the “Intensity L” one was set to 1. The proteins with an exogenous fraction in at least three out of five biological replicates were selected. We filtered out the proteins not identified in pCAF1 proteome (donor cells). For the analysis of the THP-1 proteome, the intensity value of each protein was divided by the molecular weight (MW) and transformed by log_2_. The adhesion molecules were selected based on the Gene Ontology Biological Processes (GOBP) category of cell adhesion (GO:0007155) and based on the subcellular location’s annotations retrieved from UniProt. For the analysis of pCAF1, CM-EVs and MBVs proteome, the intensity value of each protein was divided by the MW and transformed by log_2_. For the analysis of pCAF/pNF proteome, the SILAC ratio was inverted, transformed by log_2_ and normalized by subtracting the median from each column. For the analysis of the α-SMA^high^ and α-SMA^low^ pCAF proteome, LFQ intensity was transformed by log_2_, three valid values were required for at least one pCAF1 or pCAF3 subpopulation and one valid value was required for at least one pCAF4 subpopulation. Missing values were replaced from the normal distribution using the recommended setting in Perseus software and proteins with a fold change ≥ 1.5 and *P* ≤ 0.05 (two-sided Student’s T-test) in at least two out of the three pCAF lines were selected. The Z-score was calculated by row. The cell surface proteins were selected based on the subcellular location’s annotations retrieved from UniProt.

### Proteomic datasets

The proteomic data of EVs were downloaded from three publicly available datasets (51–53). For each dataset, we considered the proteins unique to each EV subpopulation and those with abundance significantly different between the two as statistically analyzed by the authors, except that for (53), for this paper we considered proteins with at least a two-fold change and *P* < 0.05. The selected proteins were matched by gene name with the proteins whose abundance was significantly different between CM-EVs and MBVs (two-sided Student’s T-test, *P* < 0.05). The Z-score was calculated by row.

### In vivo

BALB/c C.FVB-tg(Acta2-DsRed)1RK1/J mice (generated by Dr. Raghu Kalluri, MD Anderson Cancer Center, and kindly provided by Dr. Chris D Madsen, Lund University) were used for the in vivo experiments. All mouse procedures were in accordance with ethical approval from University of Glasgow under the revised Animal Act 1986 (Scientific Procedures) and the EU Directive 2010/63/EU authorized through UK Home Office Approval (Project license number 70/8645). For FACS analysis, 2.5×10^4^ 4T1 cells were resuspended in 100 μl of PBS and injected in the tail vein of 6-8-week-old RFP expressing female mice. Littermate α-SMA-RFP female mice that had not been injected with 4T1 cells were used as control. Mice were culled three weeks after the injection. Lungs were collected, minced finely and digested in pre-warmed PBS (with calcium and magnesium) with 2 mg/ml of collagenase A (Roche, Basel, Switzerland) for 1h on a rotating wheel at 37°C. The pieces of lung tissue were then pass through a 14G needle. Isolated cells were resuspended in M199 medium supplemented with 10% FBS, passed through a cell strainer (70 μm) and washed several times by centrifugation at 300 x g for 5min. Then, cells were resuspended in FACS buffer and incubated with Alexa Fluor^®^ 488 anti-CD31 ([390] 102414) and APC/Cyanine7 anti-CD45 ([30-F11] 103116) antibodies (1/100, BioLegend) and mouse TruStain FcX^TM^ (1/200, BioLegend) for 45min on ice. DAPI was used as live/dead marker. Cells were analyzed using an Attune^TM^ NxT flow cytometer and FlowJo software version 10.7.1.

For immunofluorescence analysis, 2.5×10^4^ 4T1 cells were resuspended in 100 μl of PBS and injected in the tail vein of 4-5-month-old female mice expressing or not (control mice) the RFP. Mice were culled three weeks after the injection. A small incision was made in the trachea and 1 ml of 2% low-melting point agarose was introduced slowly into the lungs through a 22G needle. Lungs were excised and fixed in 4% PFA for 2h at 4°C. Then, lungs were sliced into 300 μm thick sections by using a vibrating microtome (Campden Instruments Ltd, Loughborough, UK). Slices were permeabilized for 5h in PBS with 1% BSA, 10% normal goat serum (NGS, Sigma-Aldrich), 0.3% TX-100, and 0.05% sodium azide (VWR International, Radnor, PA), incubated overnight with anti-CD31 antibody (1/200, [2H8] MA3105 Invitrogen) and then for 3h within Alexa Fluor^®^ 647 secondary antibody (Jackson Immuno Research Labs, West Grove, PA) diluted in the same buffer. DAPI was used for nuclear staining. Slices were fixed in 4% PFA for 30min, incubated for 45min with Ce3D clearing solution and mounted with the Ce3D solution (79). Images were acquired with a Zeiss LSM 880 confocal microscope in Airyscan mode (Carl Zeiss, Plan-Apochromat 63x/1.4 Oil DIC M27 objective, zoom 1.8, z-stacks of 10-14.5 μm, 41-58 slices). Images were Airyscan processed in Zen software using default settings. Imaris software (version 9.5) was used to generate the 3D images and to calculate both the distance between RFP and Pecam1 surface, and the volume of RFP surface.

### Electron Microscopy

CM-EVs and MBVs were isolated from pCAFs as described above, fixed in 4% PFA, ultracentrifuged at 100,000 x g (4°C, 90min) and resuspended in PBS. Drops of 5 μl of CM-EV and MBV suspensions were loaded onto Carbon coated 400-mesh copper grids (Agar Scientific Ltd, Stansted, UK), which had been previously glow discharged (Quorum Q150T ES High Vacuum Unit settings 20 mA/30s). Samples were left to absorb onto carbon surfaces for 30min. Grids were floated on 100 μl droplets of PBS followed by fixation on a 50 μl droplet of 1% Glutaraldehyde (Agar Scientific Ltd) for 5min. Grids were washed with distilled water before contrast staining with Uranyl Oxalate (Merck UK) pH 7.0 (10min in the dark) followed by Methylcellulose/Uranyl Acetate (Merck UK) embedding (10min on ice in the dark). Grids were scooped up on Platinum Loops and excess fluid gently drained off leaving thin films. Grids were then left to dry before picking off and storing in a grid box. Samples were viewed on a JEOL 1200 EX TEM running at 80 kV and digital images were captured using Olympus ITEM software and a Cantega 2kx2k Camera.

### Staining and confocal microscopy of human mammary tumors

Human mammary tumors were obtained through NHS Greater Glasgow and Clyde Bio-repository. Formalin-fixed paraffin-embedded tissues were cut into 4 μm thick slices. Nine independent patient samples underwent high-pH antigen retrieval prior to immunofluorescence staining. Samples were permeabilized and blocked with 10% NGS in 1% BSA and 0.3% TX-100 for 30min. Samples were stained with unconjugated anti-TWEAR/FN14 antibody (1/75, ab109365 Abcam) for 1h in 10% NGS/1% BSA/0.3% TX-100. Samples were washed and stained with anti-CD31 Alexa Fluor^®^ 488 (1/100, [JC/70A] Abcam), anti-α-SMA-Cy3 (1/1000, [1A4] C6198 Sigma-Aldrich), goat-anti-rabbit Alexa Fluor^®^ 647 (1/200, Invitrogen, Thermo Fisher Scientific) and with Hoescht-33342 (1/5000, Sigma-Aldrich). Samples were mounted with Prolong-Glass Antifade (Invitrogen, Thermo Fisher Scientific) and allowed to cure in the dark for a minimum of 24h prior to imaging. Unstained samples were mounted as auto-fluorescence controls.

Fluorescent samples were imaged using a Zeiss 880 LSM confocal microscope (Carl Zeiss) in Lambda mode with a 32-channel spectral detector, and spectral unmixing was performed to remove as much tissue auto-fluorescence as possible. The auto-fluorescence spectrum was obtained from an unstained control, and fluorescence spectra were obtained from individual dyes (Hoescht, Alexa Fluor-488, Cy3, Alexa Fluor-647) using 405 nm, 488 nm, 561 nm and 647 nm lasers. Unbiased imaging of entire tissue sections was performed using a Plan-Apochromat 20x/0.8 M27 objective using tilescan and Z-stack modes. Tile stitching, maximum Z projection and linear unmixing was performed using Zen Black software (version 2.3 SP1), and images were visualized in Zen Blue software (version 2.3). More detailed imaging of three tissue samples was performed using a Plan-Apochromat 40x/1.3 Oil DIC M27 objective and Z-stack mode. Maximum Z projection and linear unmixing was performed as above. Image processing was performed in Fiji (ImageJ, version 1.53f51).

### Reverse transcription polymerase chain reaction (RT-qPCR)

RNA was extracted from cultured cells or cells sorted after co-culture. DNase treatment and total RNA isolation were performed using the RNeasy mini kit (Qiagen, Hilden, Germany) according to the manufacturer’s instructions. 1 μg of RNA was used to synthesize complementary DNA using the iScript kit (BioRad). DNA was diluted to 10 ng/μl and 2 μl were used in each RT-qPCR reaction with 10 μl of iTaq Universal SYBR Green Supermix (Bio-Rad Laboratories, Hercules, CA) and 400 nM of forward and reverse primers. PCR runs were performed using a QuantStudio^TM^ 3 Real-Time PCR System (Thermo Fisher Scientific). The following primers were used: TNFRSF12A: GAGAAGTTCACCACCCCCA (Fw) TGAATGAATGATGAGTGGGCGA (Rv); ACTA: GTGTGCCCCTGAAGAGCAT (Fw) GCTGGGACATTGAAAGTCTCA (Rv); TAGLN1: GGTGGAGTGGATCATCGTGC (Fw) ATGTCAGTCTTGATGACCCCA (Rv); COL1A1: TGAAGGGACACAGAGGTTTCAG (Fw) GTAGCACCATCATTTCCACGA (Rv); THY1: AGAGACTTGGATGAGGAG (Fw) CTGAGAATGCTGGAGATG (Rv); IL6: GGTACATCCTCGACGGCATCT (Fw) GTGCCTCTTTGCTGCTTTCAC (Rv); CXCL12: CTACAGATGCCCATGCCGAT (Fw) CAGCCGGGCTACAATCTGAA (Rv); 18S: AGGAATTGACGGAAGGGCAC (Fw) GGACATCTAAGGGCATCACA (Rv).

### Single-cell RNA sequencing

Data were analyzed as described in the original manuscripts (7, 16).

### Statistical analysis

Statistical analysis was performed on biologically independent replicates (N) using GraphPad Prism software version 9 (GraphPad Software Inc., San Diego, CA) and *P* was calculated as detailed in each figure legend.

## Acknowledgments

We thank Tom Gilbey, the BSU and BAIR at the Cancer Research UK Beatson Institute, NHS Greater Glasgow and Clyde Bio-repository for providing patient samples and the PRIDE Team.

## Funding

This work was funded by:

Cancer Research UK Beatson Institute grant A31287.

Cancer Research UK Glasgow Centre grant A18076.

Cancer Research UK Beatson Institute Advanced Technology Facilities A17196.

Stand Up to Cancer campaign for Cancer Research UK grant A29800 (SZ)

Cancer Research UK grant A29799 (KB).

Cancer Research UK grant A23983 (LMC).

Breast Cancer Now project grant 2019AugPR1307 (SZ).

Breast Cancer Now project grant 2019DecPR1424 (FF and LMC).

## Author contributions

Conceptualization: AS, SZ

Methodology: AS, EJK, LJN, SL, MM, NRP, FF, GK, GRB, DA, SM, MH, YK, LMC

Investigation: AS, EJK, LJN, LM, MM, NRP, YK

Visualization: AS, LJN, GK, NRP

Supervision: FM-G, CN, KB, LMC, SZ

Writing—original draft: AS

Writing—review & editing: EJK, LMC, SZ

## Competing interests

Authors declare that they have no competing interests.

## Data and materials availability

The raw files and the MaxQuant search results files have been deposited to the ProteomeXchange Consortium via the PRIDE partner repository (80) with the dataset identifier PXD034450 (proteins transferred from cancer-associated fibroblasts to endothelial cells: trans-SILAC), PXD034460 (THP-1 proteome), PXD034467 (proteomic analysis of CM-EVs and MBVs secreted by pCAFs), PXD034519 (proteomic analysis of pCAFs expressing high and low α-SMA protein levels), PXD034573 (pCAF1 proteome). pCAF/pNF proteomic dataset is available upon request.

## Notes

### Competing Interest Statement

The authors have declared no competing interest.

## References

1. Gerhart J. 1998 Warkany lecture: signaling pathways in development. Teratology. 1999;60(4):226–39.

2. Pires-daSilva A, Sommer RJ. The evolution of signalling pathways in animal development. Nat Rev Genet. 2003;4(1):39–49.

3. McCrea PD, Gu D, Balda MS. Junctional music that the nucleus hears: cell-cell contact signaling and the modulation of gene activity. Cold Spring Harb Perspect Biol. 2009;1(4):a002923.

4. Brucher BL, Jamall IS. Epistemology of the origin of cancer: a new paradigm. BMC Cancer. 2014;14:331.

5. Brucher BL, Jamall IS. Cell-cell communication in the tumor microenvironment, carcinogenesis, and anticancer treatment. Cell Physiol Biochem. 2014;34(2):213–43.

6. Santi A, Kugeratski FG, Zanivan S. Cancer Associated Fibroblasts: The Architects of Stroma Remodeling. Proteomics. 2018;18(5-6):e1700167.

7. Wu SZ, Roden DL, Wang C, Holliday H, Harvey K, Cazet AS, et al. Stromal cell diversity associated with immune evasion in human triple-negative breast cancer. EMBO J. 2020;39(19):e104063.

8. Walker RA. The complexities of breast cancer desmoplasia. Breast Cancer Res. 2001;3(3):143–5.

9. Nissen NI, Karsdal M, Willumsen N. Collagens and Cancer associated fibroblasts in the reactive stroma and its relation to Cancer biology. J Exp Clin Cancer Res. 2019;38(1):115.

10. Marsh T, Pietras K, McAllister SS. Fibroblasts as architects of cancer pathogenesis. Biochim Biophys Acta. 2013;1832(7):1070–8.

11. Kalluri R. The biology and function of fibroblasts in cancer. Nat Rev Cancer. 2016;16(9):582–98.

12. Monteran L, Erez N. The Dark Side of Fibroblasts: Cancer-Associated Fibroblasts as Mediators of Immunosuppression in the Tumor Microenvironment. Front Immunol. 2019;10:1835.

13. Sahai E, Astsaturov I, Cukierman E, DeNardo DG, Egeblad M, Evans RM, et al. A framework for advancing our understanding of cancer-associated fibroblasts. Nat Rev Cancer. 2020;20(3):174–86.

14. Shoucair I, Weber Mello F, Jabalee J, Maleki S, Garnis C. The Role of Cancer-Associated Fibroblasts and Extracellular Vesicles in Tumorigenesis. Int J Mol Sci. 2020;21(18).

15. Biffi G, Tuveson DA. Diversity and Biology of Cancer-Associated Fibroblasts. Physiol Rev. 2021;101(1):147–76.

16. Kieffer Y, Hocine HR, Gentric G, Pelon F, Bernard C, Bourachot B, et al. Single-Cell Analysis Reveals Fibroblast Clusters Linked to Immunotherapy Resistance in Cancer. Cancer Discov. 2020;10(9):1330–51.

17. Ohlund D, Handly-Santana A, Biffi G, Elyada E, Almeida AS, Ponz-Sarvise M, et al. Distinct populations of inflammatory fibroblasts and myofibroblasts in pancreatic cancer. J Exp Med. 2017;214(3):579–96.

18. Sousa CM, Biancur DE, Wang X, Halbrook CJ, Sherman MH, Zhang L, et al. Pancreatic stellate cells support tumour metabolism through autophagic alanine secretion. Nature. 2016;536(7617):479-83.

19. Yang L, Achreja A, Yeung TL, Mangala LS, Jiang D, Han C, et al. Targeting Stromal Glutamine Synthetase in Tumors Disrupts Tumor Microenvironment-Regulated Cancer Cell Growth. Cell Metab. 2016;24(5):685–700.

20. Zhao H, Yang L, Baddour J, Achreja A, Bernard V, Moss T, et al. Tumor microenvironment derived exosomes pleiotropically modulate cancer cell metabolism. Elife. 2016;5:e10250.

21. Santi A, Caselli A, Ranaldi F, Paoli P, Mugnaioni C, Michelucci E, et al. Cancer associated fibroblasts transfer lipids and proteins to cancer cells through cargo vesicles supporting tumor growth. Biochim Biophys Acta. 2015;1853(12):3211–23.

22. Toti A, Santi A, Pardella E, Nesi I, Tomasini R, Mello T, et al. Activated fibroblasts enhance cancer cell migration by microvesicles-mediated transfer of Galectin-1. J Cell Commun Signal. 2021;15(3):405–19.

23. Spees JL, Olson SD, Whitney MJ, Prockop DJ. Mitochondrial transfer between cells can rescue aerobic respiration. Proc Natl Acad Sci U S A. 2006;103(5):1283–8.

24. Ippolito L, Morandi A, Taddei ML, Parri M, Comito G, Iscaro A, et al. Cancer-associated fibroblasts promote prostate cancer malignancy via metabolic rewiring and mitochondrial transfer. Oncogene. 2019;38(27):5339–55.

25. Davis DM. Intercellular transfer of cell-surface proteins is common and can affect many stages of an immune response. Nat Rev Immunol. 2007;7(3):238–43.

26. Smyth LA, Afzali B, Tsang J, Lombardi G, Lechler RI. Intercellular transfer of MHC and immunological molecules: molecular mechanisms and biological significance. Am J Transplant. 2007;7(6):1442–9.

27. Rechavi O, Goldstein I, Kloog Y. Intercellular exchange of proteins: the immune cell habit of sharing. FEBS Lett. 2009;583(11):1792–9.

28. Rechavi O, Kalman M, Fang Y, Vernitsky H, Jacob-Hirsch J, Foster LJ, et al. Trans- SILAC: sorting out the non-cell-autonomous proteome. Nat Methods. 2010;7(11):923–7.

29. Ahmed KA, Xiang J. Mechanisms of cellular communication through intercellular protein transfer. J Cell Mol Med. 2011;15(7):1458–73.

30. Thery C, Witwer KW, Aikawa E, Alcaraz MJ, Anderson JD, Andriantsitohaina R, et al. Minimal information for studies of extracellular vesicles 2018 (MISEV2018): a position statement of the International Society for Extracellular Vesicles and update of the MISEV2014 guidelines. J Extracell Vesicles. 2018;7(1):1535750.

31. Mathieu M, Martin-Jaular L, Lavieu G, Thery C. Specificities of secretion and uptake of exosomes and other extracellular vesicles for cell-to-cell communication. Nat Cell Biol. 2019;21(1):9–17.

32. Huleihel L, Hussey GS, Naranjo JD, Zhang L, Dziki JL, Turner NJ, et al. Matrix-bound nanovesicles within ECM bioscaffolds. Sci Adv. 2016;2(6):e1600502.

33. Hussey GS, Pineda Molina C, Cramer MC, Tyurina YY, Tyurin VA, Lee YC, et al. Lipidomics and RNA sequencing reveal a novel subpopulation of nanovesicle within extracellular matrix biomaterials. Sci Adv. 2020;6(12):eaay4361.

34. Huleihel L, Bartolacci JG, Dziki JL, Vorobyov T, Arnold B, Scarritt ME, et al. Matrix-Bound Nanovesicles Recapitulate Extracellular Matrix Effects on Macrophage Phenotype. Tissue Eng Part A. 2017;23(21-22):1283–94.

35. Kay EJ, Paterson K, Riero-Domingo C, Sumpton D, Dabritz JHM, Tardito S, et al. Cancer-associated fibroblasts require proline synthesis by PYCR1 for the deposition of pro-tumorigenic extracellular matrix. Nat Metab. 2022;4(6):693–710.

36. LeBleu VS, Teng Y, O’Connell JT, Charytan D, Muller GA, Muller CA, et al. Identification of human epididymis protein-4 as a fibroblast-derived mediator of fibrosis. Nat Med. 2013;19(2):227–31.

37. Venning FA, Zornhagen KW, Wullkopf L, Sjolund J, Rodriguez-Cupello C, Kjellman P, et al. Deciphering the temporal heterogeneity of cancer-associated fibroblast subpopulations in breast cancer. J Exp Clin Cancer Res. 2021;40(1):175.

38. Rautou PE, Leroyer AS, Ramkhelawon B, Devue C, Duflaut D, Vion AC, et al. Microparticles from human atherosclerotic plaques promote endothelial ICAM-1-dependent monocyte adhesion and transendothelial migration. Circ Res. 2011;108(3):335–43.

39. Huang A, Dong J, Li S, Wang C, Ding H, Li H, et al. Exosomal transfer of vasorin expressed in hepatocellular carcinoma cells promotes migration of human umbilical vein endothelial cells. Int J Biol Sci. 2015;11(8):961–9.

40. Kannan K, Stewart RM, Bounds W, Carlsson SR, Fukuda M, Betzing KW, et al. Lysosome-associated membrane proteins h-LAMP1 (CD107a) and h-LAMP2 (CD107b) are activation-dependent cell surface glycoproteins in human peripheral blood mononuclear cells which mediate cell adhesion to vascular endothelium. Cell Immunol. 1996;171(1):10–9.

41. Sarafian V, Jadot M, Foidart JM, Letesson JJ, Van den Brule F, Castronovo V, et al. Expression of Lamp-1 and Lamp-2 and their interactions with galectin-3 in human tumor cells. Int J Cancer. 1998;75(1):105–11.

42. Dedieu S, Langlois B. LRP-1: a new modulator of cytoskeleton dynamics and adhesive complex turnover in cancer cells. Cell Adh Migr. 2008;2(2):77–80.

43. Min BK, Suk K, Lee WH. Stimulation of CD107 affects LPS-induced cytokine secretion and cellular adhesion through the ERK signaling pathway in the human macrophage-like cell line, THP-1. Cell Immunol. 2013;281(2):122–8.

44. Potere N, Del Buono MG, Mauro AG, Abbate A, Toldo S. Low Density Lipoprotein Receptor-Related Protein-1 in Cardiac Inflammation and Infarct Healing. Front Cardiovasc Med. 2019;6:51.

45. Herrera-Molina R, Valdivia A, Kong M, Alvarez A, Cardenas A, Quest AF, et al. Thy-1-interacting molecules and cellular signaling in cis and trans. Int Rev Cell Mol Biol. 2013;305:163–216.

46. Fiore VF, Ju L, Chen Y, Zhu C, Barker TH. Dynamic catch of a Thy-1-alpha5beta1+syndecan-4 trimolecular complex. Nat Commun. 2014;5:4886.

47. Singh A, Fedele C, Lu H, Nevalainen MT, Keen JH, Languino LR. Exosome-mediated Transfer of alphavbeta3 Integrin from Tumorigenic to Nontumorigenic Cells Promotes a Migratory Phenotype. Mol Cancer Res. 2016;14(11):1136–46.

48. Nakamura K, Sawada K, Kinose Y, Yoshimura A, Toda A, Nakatsuka E, et al. Exosomes Promote Ovarian Cancer Cell Invasion through Transfer of CD44 to Peritoneal Mesothelial Cells. Mol Cancer Res. 2017;15(1):78–92.

49. Shen X, Wang C, Zhu H, Wang Y, Wang X, Cheng X, et al. Exosome-mediated transfer of CD44 from high-metastatic ovarian cancer cells promotes migration and invasion of low-metastatic ovarian cancer cells. J Ovarian Res. 2021;14(1):38.

50. Kugeratski FG, Hodge K, Lilla S, McAndrews KM, Zhou X, Hwang RF, et al. Quantitative proteomics identifies the core proteome of exosomes with syntenin-1 as the highest abundant protein and a putative universal biomarker. Nat Cell Biol. 2021;23(6):631–41.

51. Xu R, Greening DW, Rai A, Ji H, Simpson RJ. Highly-purified exosomes and shed microvesicles isolated from the human colon cancer cell line LIM1863 by sequential centrifugal ultrafiltration are biochemically and functionally distinct. Methods. 2015;87:11–25.

52. Minciacchi VR, You S, Spinelli C, Morley S, Zandian M, Aspuria PJ, et al. Large oncosomes contain distinct protein cargo and represent a separate functional class of tumor-derived extracellular vesicles. Oncotarget. 2015;6(13):11327–41.

53. Kowal J, Arras G, Colombo M, Jouve M, Morath JP, Primdal-Bengtson B, et al. Proteomic comparison defines novel markers to characterize heterogeneous populations of extracellular vesicle subtypes. Proc Natl Acad Sci U S A. 2016;113(8):E968–77.

54. Sebastian A, Hum NR, Martin KA, Gilmore SF, Peran I, Byers SW, et al. Single-Cell Transcriptomic Analysis of Tumor-Derived Fibroblasts and Normal Tissue-Resident Fibroblasts Reveals Fibroblast Heterogeneity in Breast Cancer. Cancers (Basel). 2020;12(5).

55. Pedrioli G, Paganetti P. Hijacking Endocytosis and Autophagy in Extracellular Vesicle Communication: Where the Inside Meets the Outside. Front Cell Dev Biol. 2020;8:595515.

56. Gurung S, Perocheau D, Touramanidou L, Baruteau J. The exosome journey: from biogenesis to uptake and intracellular signalling. Cell Commun Signal. 2021;19(1):47.

57. Zheng J, Tan J, Miao YY, Zhang Q. Extracellular vesicles degradation pathway based autophagy lysosome pathway. Am J Transl Res. 2019;11(3):1170–83.

58. Kaushik S, Pickup MW, Weaver VM. From transformation to metastasis: deconstructing the extracellular matrix in breast cancer. Cancer Metastasis Rev. 2016;35(4):655–67.

59. Reid SE, Kay EJ, Neilson LJ, Henze AT, Serneels J, McGhee EJ, et al. Tumor matrix stiffness promotes metastatic cancer cell interaction with the endothelium. EMBO J. 2017;36(16):2373–89.

60. Winkler J, Abisoye-Ogunniyan A, Metcalf KJ, Werb Z. Concepts of extracellular matrix remodelling in tumour progression and metastasis. Nat Commun. 2020;11(1):5120.

61. Albacete-Albacete L, Navarro-Lerida I, Lopez JA, Martin-Padura I, Astudillo AM, Ferrarini A, et al. ECM deposition is driven by caveolin-1-dependent regulation of exosomal biogenesis and cargo sorting. J Cell Biol. 2020;219(11).

62. Nawaz M, Shah N, Zanetti BR, Maugeri M, Silvestre RN, Fatima F, et al. Extracellular Vesicles and Matrix Remodeling Enzymes: The Emerging Roles in Extracellular Matrix Remodeling, Progression of Diseases and Tissue Repair. Cells. 2018;7(10).

63. Becker A, Thakur BK, Weiss JM, Kim HS, Peinado H, Lyden D. Extracellular Vesicles in Cancer: Cell-to-Cell Mediators of Metastasis. Cancer Cell. 2016;30(6):836–48.

64. Commisso C, Davidson SM, Soydaner-Azeloglu RG, Parker SJ, Kamphorst JJ, Hackett S, et al. Macropinocytosis of protein is an amino acid supply route in Ras-transformed cells. Nature. 2013;497(7451):633–7.

65. Hosios AM, Hecht VC, Danai LV, Johnson MO, Rathmell JC, Steinhauser ML, et al. Amino Acids Rather than Glucose Account for the Majority of Cell Mass in Proliferating Mammalian Cells. Dev Cell. 2016;36(5):540–9.

66. Lin XP, Mintern JD, Gleeson PA. Macropinocytosis in Different Cell Types: Similarities and Differences. Membranes (Basel). 2020;10(8).

67. Li X, Sun X, Carmeliet P. Hallmarks of Endothelial Cell Metabolism in Health and Disease. Cell Metab. 2019;30(3):414–33.

68. Comito G, Ippolito L, Chiarugi P, Cirri P. Nutritional Exchanges Within Tumor Microenvironment: Impact for Cancer Aggressiveness. Front Oncol. 2020;10:396.

69. Bergers G, Song S. The role of pericytes in blood-vessel formation and maintenance. Neuro Oncol. 2005;7(4):452–64.

70. Latif N, Sarathchandra P, Chester AH, Yacoub MH. Expression of smooth muscle cell markers and co-activators in calcified aortic valves. Eur Heart J. 2015;36(21):1335–45.

71. Alvarez-Castelao B, Schanzenbacher CT, Hanus C, Glock C, Tom Dieck S, Dorrbaum AR, et al. Cell-type-specific metabolic labeling of nascent proteomes in vivo. Nat Biotechnol. 2017;35(12):1196–201.

72. Kojima Y, Acar A, Eaton EN, Mellody KT, Scheel C, Ben-Porath I, et al. Autocrine TGF-beta and stromal cell-derived factor-1 (SDF-1) signaling drives the evolution of tumor-promoting mammary stromal myofibroblasts. Proc Natl Acad Sci U S A. 2010;107(46):20009–14.

73. Rappsilber J, Ishihama Y, Mann M. Stop and go extraction tips for matrix-assisted laser desorption/ionization, nanoelectrospray, and LC/MS sample pretreatment in proteomics. Anal Chem. 2003;75(3):663–70.

74. Wisniewski JR, Zougman A, Nagaraj N, Mann M. Universal sample preparation method for proteome analysis. Nat Methods. 2009;6(5):359–62.

75. Kugeratski FG, Atkinson SJ, Neilson LJ, Lilla S, Knight JRP, Serneels J, et al. Hypoxic cancer-associated fibroblasts increase NCBP2-AS2/HIAR to promote endothelial sprouting through enhanced VEGF signaling. Sci Signal. 2019;12(567).

76. Shevchenko A, Tomas H, Havlis J, Olsen JV, Mann M. In-gel digestion for mass spectrometric characterization of proteins and proteomes. Nat Protoc. 2006;1(6):2856–60.

77. Tyanova S, Temu T, Cox J. The MaxQuant computational platform for mass spectrometry-based shotgun proteomics. Nat Protoc. 2016;11(12):2301–19.

78. Tyanova S, Temu T, Sinitcyn P, Carlson A, Hein MY, Geiger T, et al. The Perseus computational platform for comprehensive analysis of (prote)omics data. Nat Methods. 2016;13(9):731–40.

79. Li W, Germain RN, Gerner MY. High-dimensional cell-level analysis of tissues with Ce3D multiplex volume imaging. Nat Protoc. 2019;14(6):1708–33.

80. Vizcaino JA, Cote RG, Csordas A, Dianes JA, Fabregat A, Foster JM, et al. The PRoteomics IDEntifications (PRIDE) database and associated tools: status in 2013. Nucleic Acids Res. 2013;41(Database issue):D1063–9.

